# Identification of lectin receptors for conserved SARS-CoV-2 glycosylation sites

**DOI:** 10.1101/2021.04.01.438087

**Authors:** David Hoffmann, Stefan Mereiter, Yoo Jin Oh, Vanessa Monteil, Rong Zhu, Daniel Canena, Lisa Hain, Elisabeth Laurent, Clemens Grünwald-Gruber, Maria Novatchkova, Melita Ticevic, Antoine Chabloz, Gerald Wirnsberger, Astrid Hagelkruys, Friedrich Altmann, Lukas Mach, Johannes Stadlmann, Chris Oostenbrink, Ali Mirazimi, Peter Hinterdorfer, Josef M. Penninger

**Author notes:** These authors contributed equally to this work (David Hoffmann, Stefan Mereiter).

## Abstract

New SARS-CoV-2 variants are continuously emerging with critical implications for therapies or vaccinations. All 22 N-glycan sites of SARS-CoV-2 Spike remain highly conserved among the variants B.1.1.7, 501Y.V2 and P.1, opening an avenue for robust therapeutic intervention. Here we used a comprehensive library of mammalian carbohydrate-binding proteins (lectins) to probe critical sugar residues on the full-length trimeric Spike and the receptor binding domain (RBD) of SARS-CoV-2. Two lectins, Clec4g and CD209c, were identified to strongly bind to Spike. Clec4g and CD209c binding to Spike was dissected and visualized in real time and at single molecule resolution using atomic force microscopy. 3D modelling showed that both lectins can bind to a glycan within the RBD-ACE2 interface and thus interferes with Spike binding to cell surfaces. Importantly, Clec4g and CD209c significantly reduced SARS-CoV-2 infections. These data report the first extensive map and 3D structural modelling of lectin-Spike interactions and uncovers candidate receptors involved in Spike binding and SARS-CoV-2 infections. The capacity of CLEC4G and mCD209c lectins to block SARS-CoV-2 viral entry holds promise for pan-variant therapeutic interventions.

## Introduction

COVID-19 caused by SARS-CoV-2 infections has triggered a pandemic massively disrupting health care, social and economic life. SARS-CoV-2 main entry route into target cells is mediated by the viral Spike protein, which binds to angiotensin converting enzyme 2 (ACE2) expressed on host cells (Monteil et al., 2020). The Spike protein is divided into two subunits, S1 and S2. The S1 subunits comprises the receptor binding domain (RBD) which confers ACE2 binding activity. The S2 subunit mediates virus fusion with the cell wall following proteolytic cleavage (Hoffmann et al., 2020; Shang et al., 2020; Walls et al., 2020). Cryo-electron microscopy studies have shown that the Spike protein forms a highly flexible homotrimer containing 22 *N*-glycosylation sites each, 18 of which are conserved with the closely related SARS-CoV which caused the 2002/03 SARS epidemic (Ke et al., 2020; Walls et al., 2020). Point mutations removing glycosylation sites of the SARS-CoV-2 Spike protein were found to yield less infectious pseudo-typed viruses (Li et al., 2020). As Spike and RBD glycosylation affect ACE2 binding and SARS-CoV-2 infections, targeting virus specific glycosylation could be a novel means for therapeutic intervention.

Glycosylation of viral proteins ensures proper folding and shields antigenic viral epitopes from immune recognition (Watanabe et al., 2020b; Watanabe et al., 2019). To create this glycan shield, the virus hijacks the host glycosylation machinery and thereby ensures the presentation of self-associated glycan epitopes. Apart from shielding epitopes from antibody recognition, glycans can be ligands for lectin receptors. For instance, mannose-specific mammalian lectins, like DC-SIGN (CD209) or its homolog L-SIGN (CD299), are well known to bind to viruses like HIV-1 and also SARS-CoV (Van Breedam et al., 2014). Lectin receptors are often expressed on immune and endothelial cells and serve as pattern recognition receptors involved in virus internalization and transmission (Osorio and Reis e Sousa, 2011). Recent studies have characterized the recognition of the SARS-CoV-2 Spike by previously known virus-binding lectins, such as DC-SIGN, L-SIGN, MGL and MR (Gao et al., 2020). Given that SARS-CoV-2 relies less on oligo-mannose-type glycosylation, as compared to for instance HIV-1, and displays more complex-type glycosylation, it is unknown if additional lectin receptors are capable of binding the Spike protein and whether such interactions might have functional relevance in SARS-CoV-2 infections.

## Results

### Preparation of the first near genome-wide lectin library to screen for novel binders of Spike glycosylation

To systematically identify lectins that bind to the trimeric Spike protein and RBD of SARS-CoV-2, we searched for all annotated carbohydrate recognition domains (CRDs) of mouse C-type lectins, Galectins and Siglecs. Of 168 annotated CRDs, we were able to clone, express and purify 143 lectin-CRDs as IgG2a-Fc fusion proteins from human HEK293F cells (Fig. 1A, table S1). The resulting dimeric lectin-Fc fusion proteins (hereafter referred to as lectins) showed a high degree of purity (Fig. 1B). This collection of lectins is, to our knowledge, the first comprehensive library of mammalian CRDs.

**Figure 1.**
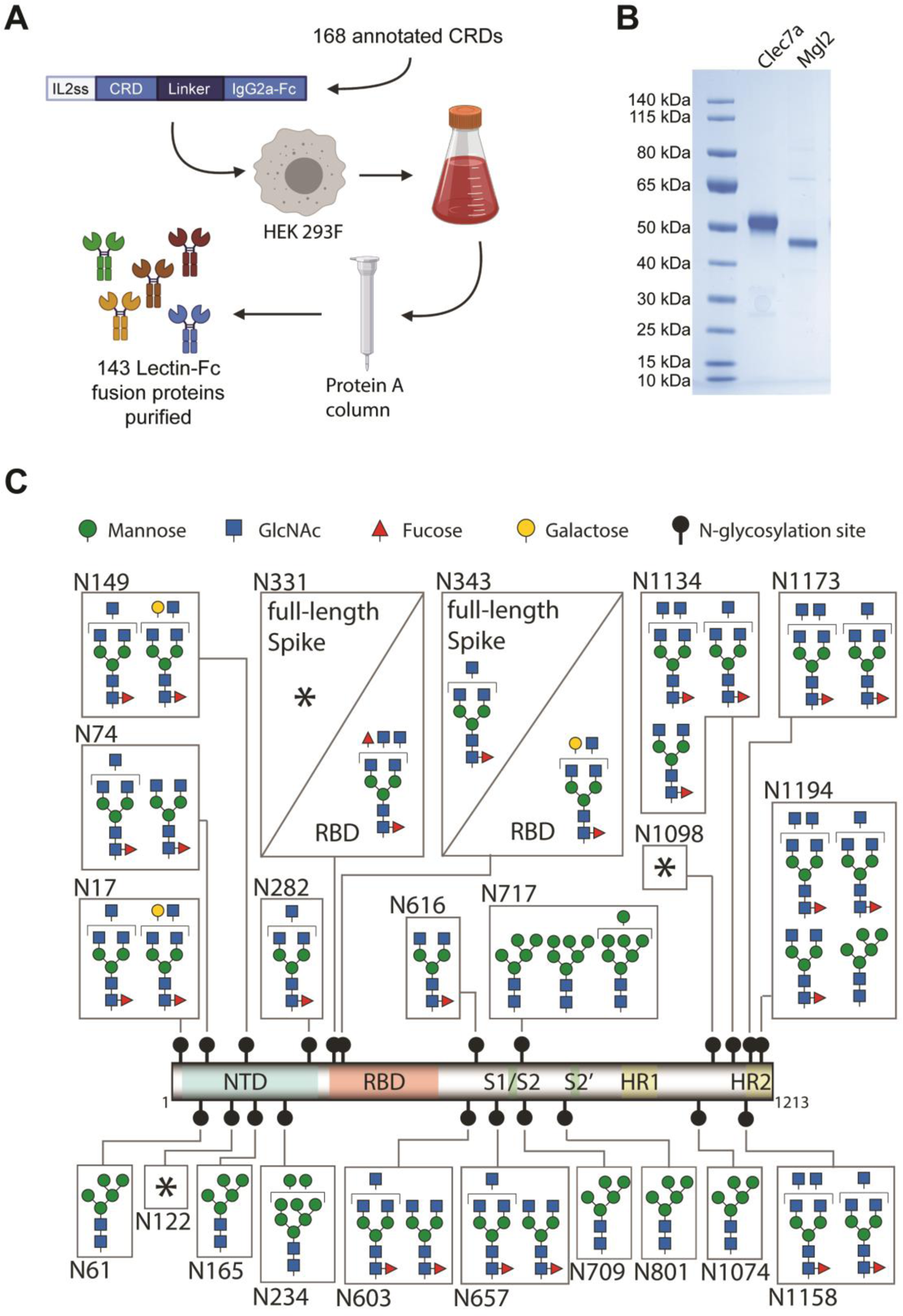
Lectin library and SARS-CoV-2 Spike and RBD glycosylation. (**A**) Schematic overview of cloning, expression and purification of 143 carbohydrate recognition domain (CRD) – mouse IgG2a-Fc fusion proteins, from 168 annotated murine CRD containing proteins. The constructs were expressed in HEK293F cells and secreted Fc-fusion proteins were purified using protein A columns. See table S1 for full list of expressed CRDs. (**B**) Exemplified SDS-PAGE of purified Clec7a and Mgl2 stained with Coomassie blue. (**C**) Glycosylation map of the SARS-CoV-2 Spike and RBD. The most prominent glycan structures are represented for each site, with at least 15% relative abundance. **\*** marks highly variable glycosylation sites in which no single glycan structure accounted for >15% relative abundance. The different monosaccharides are indicated using standardized nomenclature. NTD, n-terminal domain; RBD, receptor binding domain; S1/S2 and S2’, proteolytic cleavage sites; HR1 and HR2, α-helical heptad repeat domains 1 and 2; GlcNAc, N-acetylglucosamine.

**Table 1.** Values computed for surface plasmon resonance (SPR) and atomic force microscopy (AFM). SPR. Kinetic association (k_a,1_), kinetic disssociation (k_d,1_), and equlibrium dissociation (K_d,1_) constants of the first binding step fitted from the bivalent analyte model, assuming two-step binding and dissociation of the lectins to adjacent immobilized Spike trimer binding sites under spontaneous thermodynamic energy barriers; no reasonable fit was obtained with the simple 1:1 binding model (data not shown). AFM. Kinetic off-rate constants (*k_off_*) and lengths of dissociation paths (x_β_) of single lectin bonds, originating from force-induced unbinding in single molecule force spectroscopy (SMFS) experiments and computed using Evans’s (Bell, 1978; Evans and Ritchie, 1997) model, assuming a sharp single dissociation energy barrier.

|  | SPR |  |  | AFM |  |
| --- | --- | --- | --- | --- | --- |
| | $k_{a,1}$<br>[M<sup>-1</sup>s<sup>-1</sup>] | $k_{d,1}$<br>[s<sup>-1</sup>] | $K_{d,1}$<br>[M] | $k_{off}$<br>[s<sup>-1</sup>] | $x_{\beta}$<br>[nm] |
| **Clec4g** | $6,17 \times 10^4$ | 0,0997 | $1,62 \times 10^{-6}$ | $0.037 \pm 0.007$ | $0.55 \pm 0.02$ |
| **CD209c** | $1,61 \times 10^4$ | 0,0159 | $0,988 \times 10^{-6}$ | $0.041 \pm 0.025$ | $0.76 \pm 0.03$ |
| **hCLEC4G** | $7,77 \times 10^4$ | 0,0201 | $0,259 \times 10^{-6}$ | $0.007 \pm 0.0004$ | $0.64 \pm 0.55$ |
| **hCD209** | $1,32 \times 10^4$ | 0,0316 | $2,39 \times 10^{-6}$ | $0.008 \pm 0.001$ | $0.73 \pm 0.14$ |

We next recombinantly expressed monomeric RBD and full-length trimeric Spike protein (hereafter referred to as Spike protein) in human HEK293-6E cells. Using mass spectrometry, we characterized all 22 *N*-glycosylation sites on the full-length Spike protein and 2 *N-*glycosylation sites on the RBD (Fig. 1C and table S2). Most of the identified structures were in accordance with previous studies using full-length Spike (Watanabe et al., 2020a) with the exception of N331, N603 and N1194, which presented a higher structural variability of the glycan branches (Fig. 1C and table S2). Importantly, the *N*-glycan sites of Spike are highly conserved among the sequenced SARS-CoV-2 viruses including the emerging variants B.1.1.7, 501Y.V2 and P.1 (Fig S1A, B). The detected *N*-glycan species ranged from poorly processed oligo-mannose structures to highly processed multi-antennary complex *N*-glycans in a site-dependent manner. This entailed also a large variety of terminal glycan epitopes, which could act as ligands for lectins. Notably, the two glycosylation sites N331 and N343 located in the RBD carried more extended glycans, including sialylated and di-fucosylated structures, when expressed as an independent construct as opposed to the full-length Spike protein (Fig. 1C and table S2). These data underline the complex glycosylation of Spike and reveal that *N*-glycosylation of the RBD within the 3D context of full-length trimeric Spike is different from *N*-glycosylation of the RBD expressed as minimal ACE2 binding domain.

### CD209c and Clec4g are novel high affinity binders of SARS-CoV-2 Spike

We evaluated the reactivity of our murine lectin library against the trimeric Spike and monomeric RBD of SARS-CoV-2 using an ELISA assay (Fig. S2A). This screen revealed that CD209c (SIGNR2), Clec4g (LSECtin), and Reg1 exhibited pronounced binding to Spike, whereas Mgl2 and Asgr1 displayed elevated binding to the RBD (Fig. 2A, B and table S3). Further, we investigated the reactivity of the lectin library against human recombinant soluble ACE2 (hrsACE2); none of the lectins bound to hrsACE2 (Fig. S2B). We excluded Reg1 from further studies due to inconsistent ELISA results, likely due to protein instability. Asgr1 was excluded because it bound only to RBD but not to the Spike trimer, in accordance to the differences in glycosylation of glycosites N331 and N343 between Spike and RBD (Fig. 1C). This highlights the importance of using a full-length trimeric Spike protein for functional studies. To confirm that the observed interactions were independent of protein conformation, Spike was denatured prior to the ELISA assay; binding of CD209c and Clec4g to the unfolded Spike remained unaltered (Fig. 2C). Importantly, enzymatic removal of *N*-glycans by PNGase F treatment reduced the binding of CD209c, Clec4g, and Mgl2 towards Spike (Fig. 2D and Fig. S2C), confirming *N*-glycans as ligands. Binding of ACE2, which relies on protein-protein interactions, was completely abrogated when Spike was denatured (Fig. 2C). These data identify lectins that have the potential to bind to the RBD and trimeric Spike of SARS-CoV-2.

**Figure 2.**
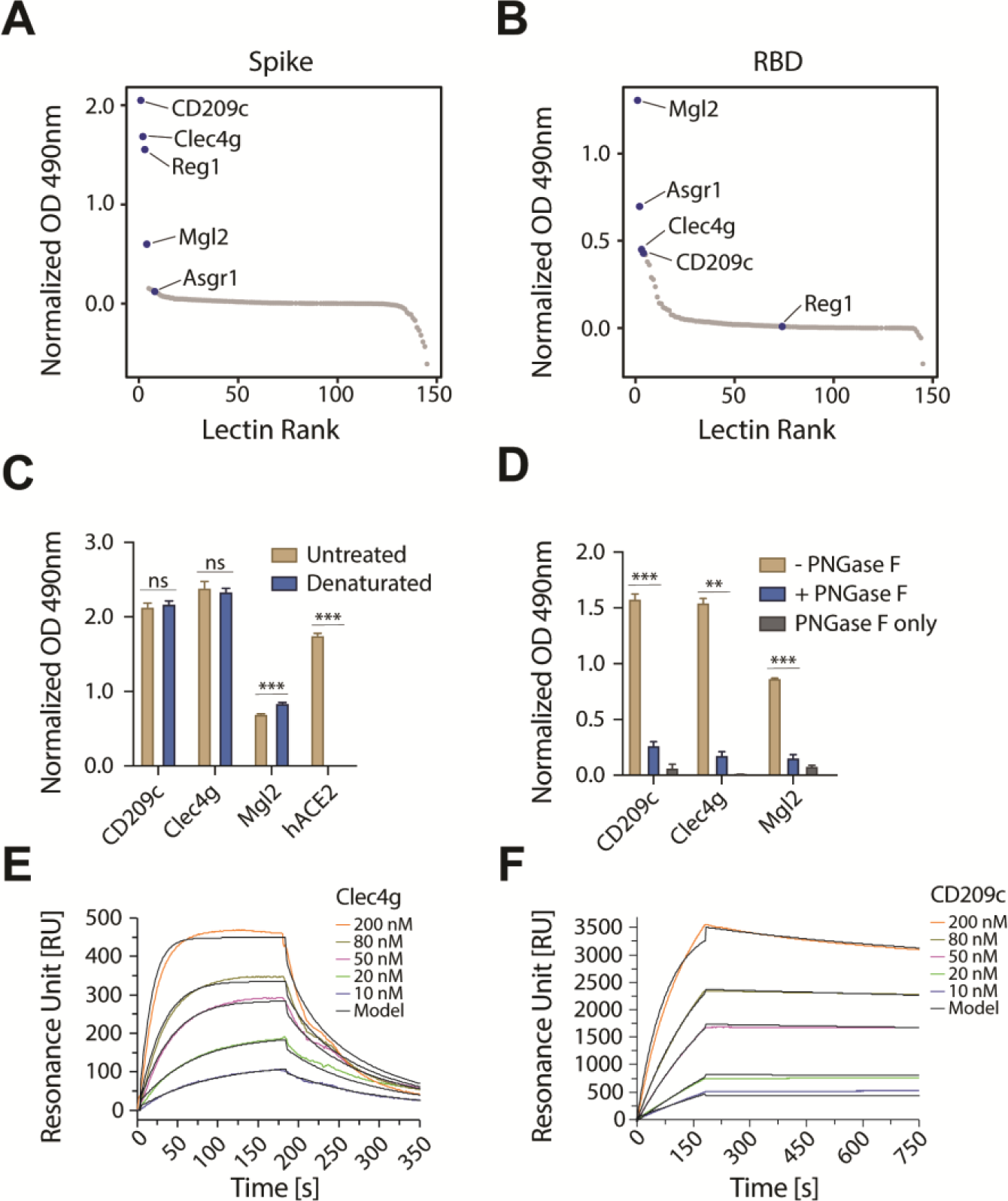
Identification of lectins that bind to Spike and RBD of SARS-CoV-2. (**A**) and (**B**) ELISA screen of the lectin-Fc library against full-length trimeric SARS-CoV-2 Spike (A) or monomeric RBD (B). Results are shown as mean OD values of 2 replicates normalized against a BSA control and ranked by value. Lectin-Fc fusion proteins with a normalized OD > 0.5 in either (A) or (B) are indicated in both panels. See table S2 for primary ELISA data. (**C**) Lectin-Fc and human ACE2-mIgG1 Fc-fusion protein (hACE2) binding to untreated or heat-denatured full-length SARS-CoV-2 Spike by ELISA. hACE2-mIgG1 was used as control for complete denaturation of Spike protein. Results are shown as mean OD values ± SD normalized to the BSA control (N=3). (**D**) Lectin-Fc binding to full-length SARS-CoV-2 Spike with or without de-*N*-glycosylation by PNGase F. “PNGase F only” denotes wells that were not coated with the Spike protein. Results are shown as mean OD values ± SD normalized to BSA controls (N=3). (**E**) and (**F**) Surface plasmon resonance (SPR) analysis with immobilized full-length trimeric Spike, probed with various concentrations of Clec4g-Fc (E) and CD209c-Fc (F). See Table 1 for kinetics values. (C) t-test with Holm-Sidak correction for multiple comparisons. (D) One-way ANOVA with Tukey’s multiple comparisons; *P<0.05; **P<0.01; ***P<0.001; ns: not significant.

Based on the robust *N*-glycan dependent Spike binding, we focused our further studies on CD209c and Clec4g. We first used surface plasmon resonance (SPR) to determine the kinetic and equilibrium binding constants of these lectins to the trimeric Spike. The resulting experimental binding curves were fitted to the “bivalent analyte model” (Traxler et al., 2017) which assumes two-step binding of the lectin dimers to adjacent immobilized Spike trimer binding sites (Fig. 2E, F). From these fits, we computed the kinetic association (k_a,1_), kinetic dissociation (k_d,1_), and equilibrium dissociation (K_d,1_) binding constants of single lectin bonds (Table 1). The equilibrium dissociations (K_d,1_) values were 1.6 μM and 1.0 μM for Clec4g and CD209c, respectively.

### Multiple CD209c and Clec4g molecules bind simultaneously to SARS-CoV-2 Spike and form compact complexes

To study Spike binding of these two lectins at the single-molecule level, we used atomic force microscopy (AFM) and performed single molecule force spectroscopy (SMFS) experiments. To this end, we coupled trimeric Spike to the tip of the AFM cantilever and performed single-molecule force measurements (Hinterdorfer et al., 1996), by moving the Spike trimer-coupled tip towards the surface-bound lectins to allow for bond formations (Fig. 3A). Unbinding was accomplished by pulling on the bonds, which resulted in characteristic downward deflection signals of the cantilever, whenever a bond was ruptured (Fig. 3B). The magnitude of these vertical jumps reflects the unbinding forces, which were of typical strengths for specific molecular interactions (Rankl et al., 2008). Using this method (Rankl et al., 2008; Zhu et al., 2010), we quantified unbinding forces (Fig. 3C) and calculated the binding probability and the number of bond ruptures between CD209c or Clec4g and trimeric Spike (Fig. 3D). Both lectins showed a very high binding probability and could establish up to 3 strong bonds with accumulating interaction force strengths reaching 150 pN in total with trimeric Spike (Fig. 3C), with the preference of single and dual bonds (Fig. 3C, 3D, Fig. S3A, B, Table 1). Of note, multi-bond formation leads to stable complex formation, in which the number of formed bonds enhances the overall interaction strength and dynamic stability of the complexes. To assess dynamic interactions between single molecules of trimeric Spike and the lectins in real time we used high-speed AFM (Kodera et al., 2010; Preiner et al., 2014). Addition of Clec4g and CD209c led to a volume increase of the lectin/Spike complex in comparison to the trimeric Spike alone; based on the volumes we could calculate that on average 3.2 molecules of Clec4g and 5.2 molecules of CD209c were bound to one Spike trimer (Fig. 3E, Fig. S3C-E and movies S1-4). These data show, in real-time, at single molecule resolution, that mouse Clec4g and CD209c can directly associate with trimeric Spike.

**Figure 3.**
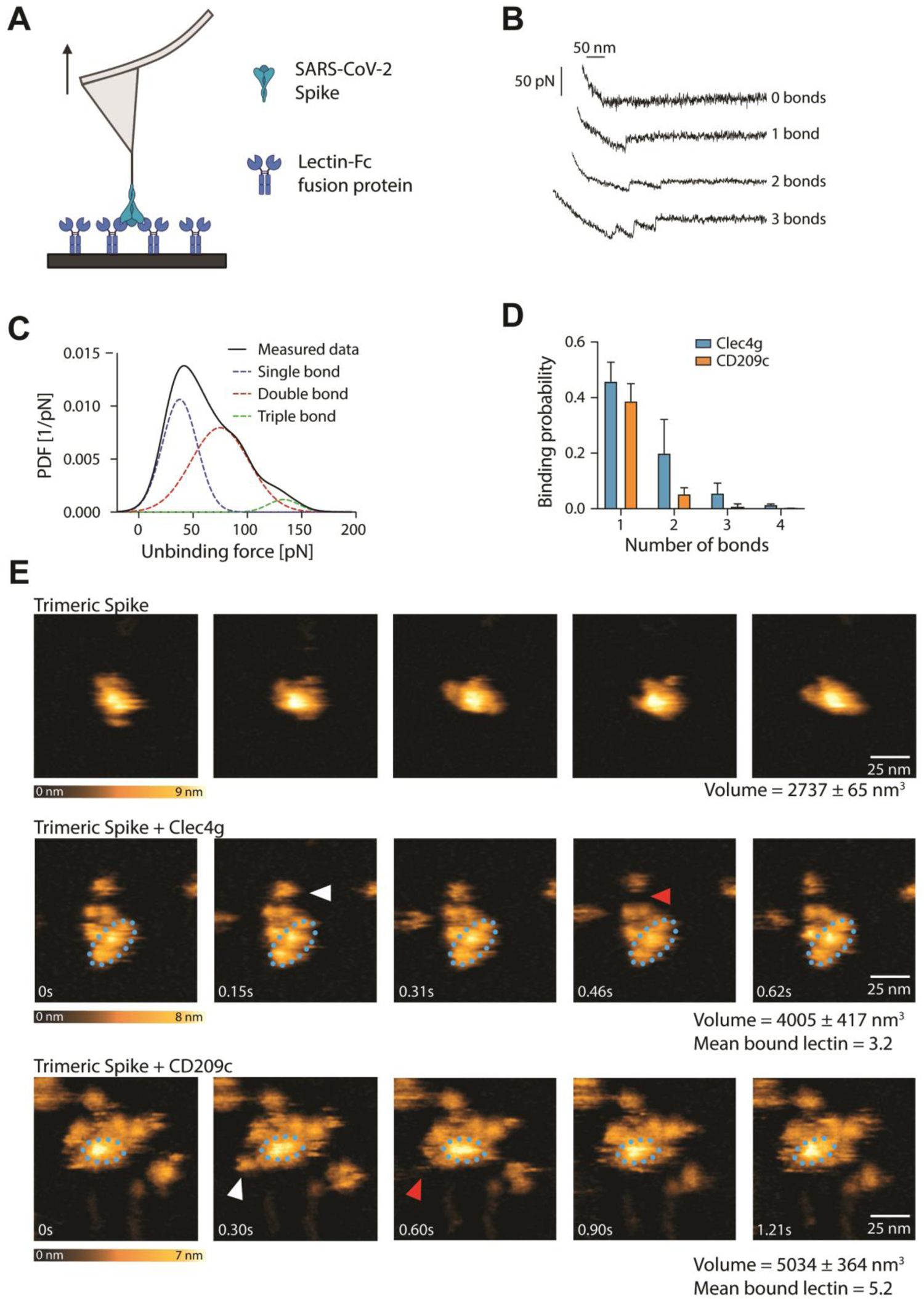
Single molecule, real time imaging of lectin-Spike binding. (**A**) Schematic overview of single molecule force spectroscopy (SMFS) experiments using full-length trimeric Spike coupled to an atomic force microscopy (AFM) cantilever tip and surface coated murine Clec4g-Fc or CD209c-Fc. Arrow indicates pulling of cantilever. (**B**) Representative force traces showing sequential bond ruptures in the SMFS experiments. Measured forces are shown in pico-Newtons (pN). (**C**) Experimental probability density function (PDF) of unbinding forces (in pN) determined by SMFS (black line, measured data). The three distinct maxima fitted by a multi-Gaussian function reveal rupture of a single bond (blue dotted line), or simultaneous rupture of 2 (red dotted line) and 3 (green dotted line) bonds, respectively. (**D**) SMFS-determined binding probability for the binding of trimeric Spike to Clec4g and CD209c. Data are shown as mean binding probability ± SD of single, double, triple or quadruple bonds (N=2). (**E**) High speed AFM of single trimeric Spike visualizing the real-time interaction dynamics with lectins. Top panel shows 5 frames of trimeric Spike alone imaged on mica. Middle and bottom panels show 5 sequential frames of trimeric Spike/Clec4g and trimeric Spike/CD209c complexes, acquired at a rate of 153.6 and 303 ms/frame, respectively. Association and dissociation events between lectin and Spike are indicated by white and red arrows, respectively. The blue dotted ellipses display the core of the complexes showing low conformational mobility. Color schemes indicate height of the molecules in nanometers (nm). Volumes of single trimeric Spike, trimeric Spike/Clec4g and trimeric Spike/CD209c complexes are indicated, as well as numbers of lectins bound to trimeric Spike, averaged over the experimental recording period.

### The human lectins CD209 and CLEC4G are high affinity receptors for SARS-CoV-2 Spike

Having characterized binding of murine CD209 and Clec4g to Spike, we next assessed whether their closest human homologues, namely human CD209 (hCD209), and human CD299 (hCD299) for murine CD209c, and human CLEC4G (hCLEC4G) for mouse Clec4g, can also bind to full-length trimeric Spike of SARS-CoV-2. hCD209, hCD299, and hCLEC4G indeed exhibited binding to Spike (Fig. 4A), demonstrating conserved substrate specificities. The binding of these human lectins was again independent of Spike folding and abrogated by *N*-glycan removal (Fig. 4A, B). SPR measurements of hCLEC4G and hCD209 bond formation to Spike showed equilibrium dissociation (K_d,1_) values of 0.3 µM and 2.4 μM, respectively, in which the high affinity of hCLEC4G is mainly contributed by its rapid kinetic association rate constant (Fig. 4C, D and Table 1). In addition, hCD209 and hCLEC4G showed a high binding probability by AFM with the formation of up to 3 bonds per trimeric Spike (Fig. S4A-C). When we monitored the dynamic interactions of the lectins with the Spike using high speed AFM, we observed binding of - on average - 3.5 hCLEC4G and 3.6 hCD209 molecules per trimeric Spike (Fig. 4E and Fig. S4D-F). In summary, our data using ELISA, SMFS, surface plasmon resonance, and high-speed atomic force microscopy show that the human lectins CLEC4G and CD209 can bind to trimeric Spike of SARS-CoV-2, in which the overall interaction strength and dynamic stability leads to compact complex formation.

**Figure 4.**
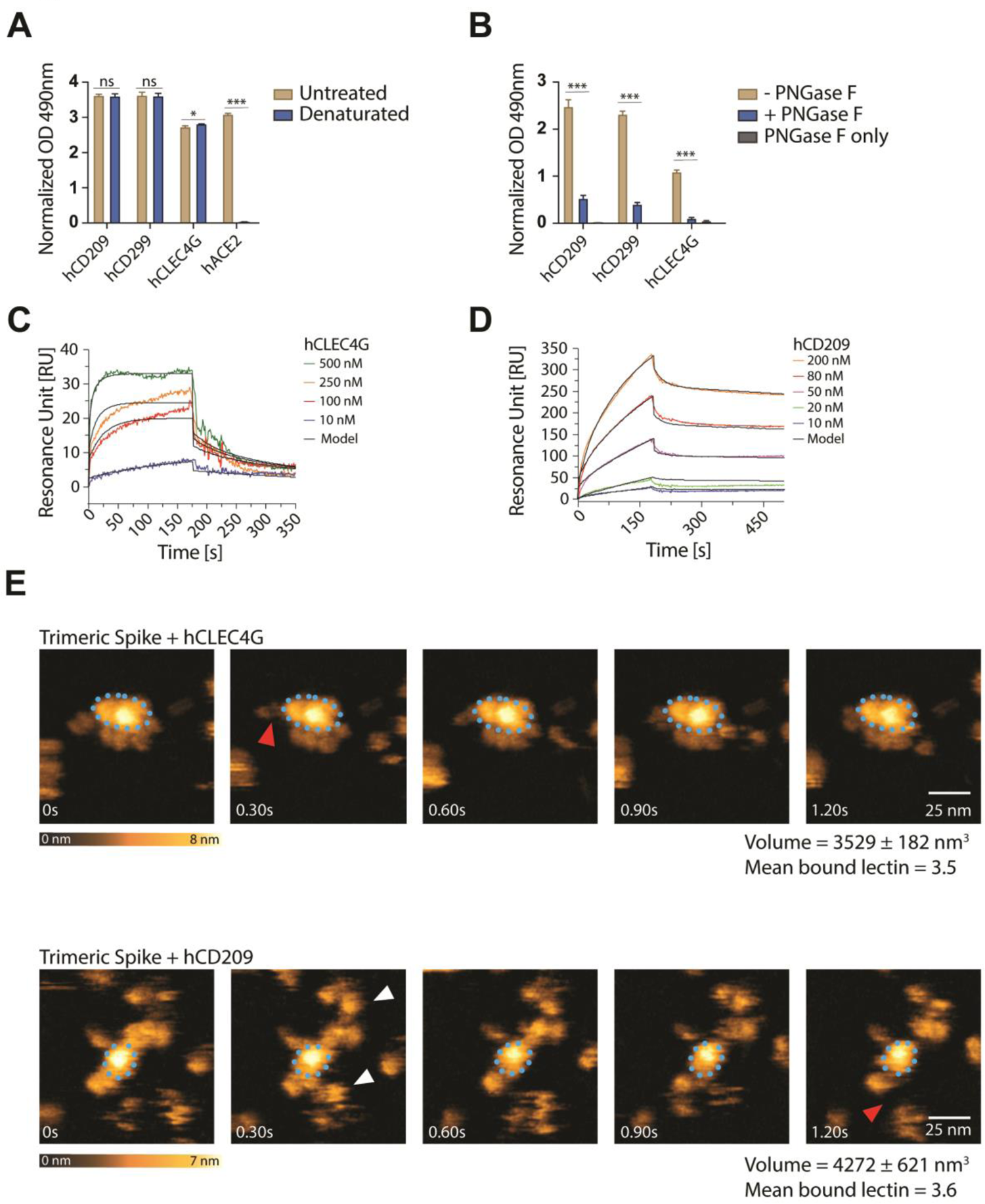
Characterization of human lectin-Spike interactions. (**A**) ELISA analyses of human lectin-hIgG1 Fc-fusion protein (hCLEC4g, hCD209, hCD299) binding to untreated or heat-denatured full-length SARS-CoV-2 Spike. A human ACE2-hIgG1-Fc fusion protein (hACE2) was used to control for the complete denaturation of Spike. Results are shown as mean OD values ± SD normalized to a BSA control (N=3). (**B**) hCLEC4g, hCD209, and hCD299 binding to full-length SARS-CoV-2 Spike with or without de-*N*-glycosylation by PNGase F. “PNGase F only” denotes wells that were not coated with the Spike protein. Results are shown as mean OD values ± SD normalized against the BSA control (N=3). (**C**) and (**D**) SPR analysis with immobilized full-length trimeric Spike, probed with various concentrations of hCLEC4G (C) and hCD209 (D). See Table 1 for kinetics values. (**E**) High speed AFM of single trimeric Spike visualizing the real-time interaction dynamics with lectins. Top and bottom panels show 5 sequential frames of trimeric Spike/hCLEC4G and trimeric Spike/hCD209 complexes, acquired at a rate of 303 ms/frame. Association and dissociation events between hCLEC4G or hCD209 and Spike are indicated by white and red arrows, respectively. The blue dotted ellipses display the core of the complexes showing low conformational mobility. Color schemes indicate height of the molecules in nanometers (nm). Volumes of single trimeric Spike and trimeric Spike/Clec4g and trimeric Spike/CD209c complexes are indicated, as well as numbers of lectins bound to trimeric Spike, averaged over the experimental recording period. (A) t-test with Holm-Sidak correction for multiple comparisons. (B) One-way ANOVA with Tukey’s multiple comparisons; *P<0.05; ***P<0.001; ns: not significant.

### CLEC4G sterically interferes with Spike/ACE2 interaction

We next 3D modelled binding of hCLEC4G and hCD209 to the candidate glycosylation sites present on Spike and how such attachment might relate to the binding of the trimeric Spike protein to its receptor ACE2. hCD209 is known to bind with high affinity to oligo-mannose structures (Guo et al., 2004). The N234 glycosylation site is the only site within the Spike that carries exclusively oligo-mannose glycans with up to 9 mannose residues (Fig. 1C, table S2). 3D modelling revealed that the oligo-mannose glycans on N234 are accessible for hCD209 binding on all 3 monomers comprising the trimeric Spike (Fig. 5 and Fig. S5A, B). Superimposition of ACE2 interacting with the RBD and hCD209 binding to N234, showed that the hCD209 binding occurs at the lateral interface of Spike, distant from the RBD (Fig. 5)

**Figure 5.**
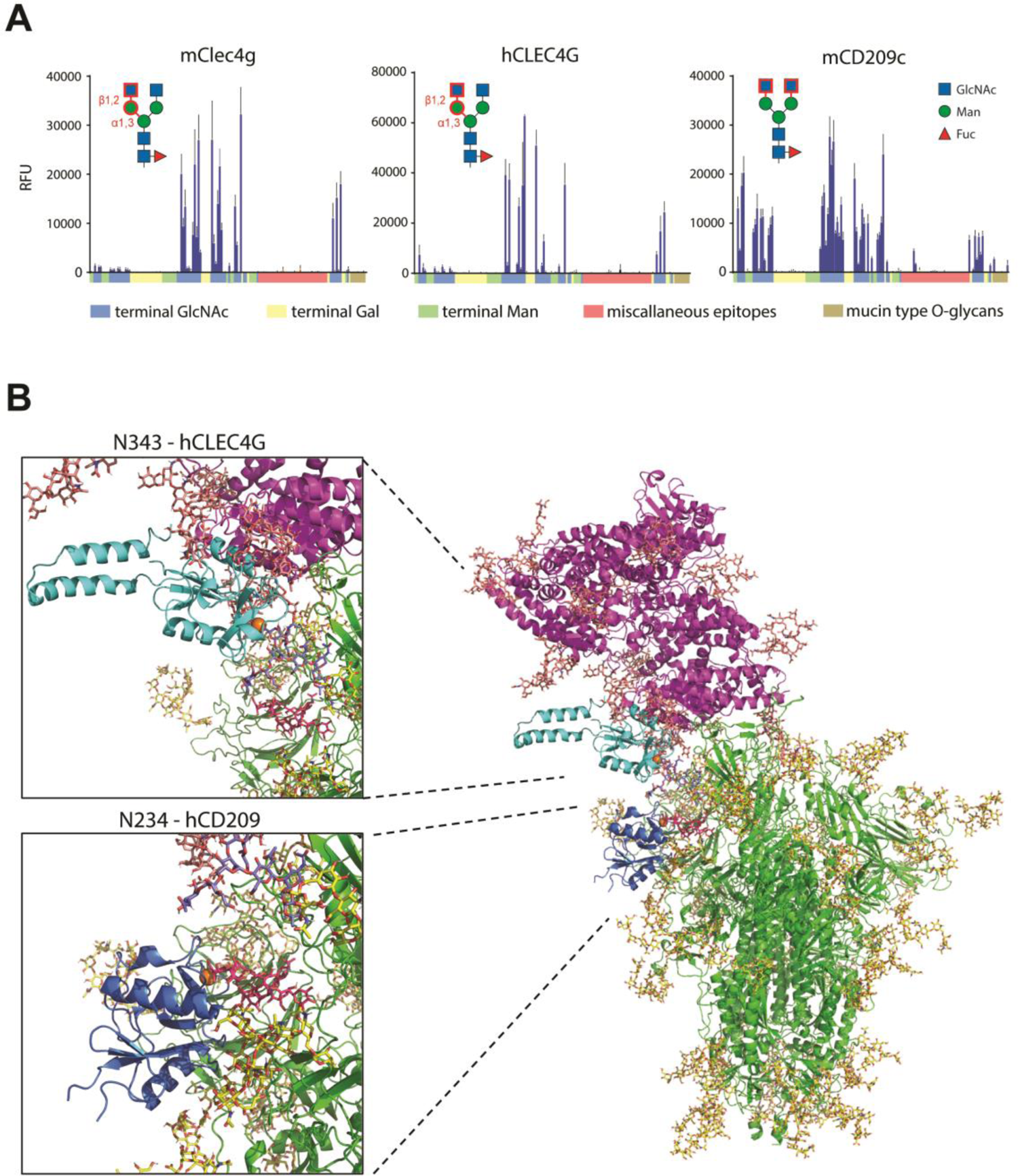
Glycan ligand characterization and structural modelling. (**A**) Glycan microarray analyses of mClec4g, mCD209c and hCLEC4G. Terminal GlcNAc, Gal or Man are colour-coded along the X-axis. Miscellaneous epitopes could not unambiguously be grouped to either terminal GlcNAc, Gal or Man. The identified main binding epitope of each lectin is shown in red. Data represent the average and standard deviation of 3 independent technical replicates. (**B**) 3D structural modelling of glycosylated trimeric Spike (green with glycans in yellow) interacting with glycosylated human ACE2 (purple with glycans in salmon). The CRD of hCLEC4G (cyan with Ca^2+^ in orange) was modelled onto Spike monomer 3 glycan site N343 (complex type glycan with terminal GlcNAc in purple-blue) and the CRD of hCD209 (dark blue with Ca^2+^ in orange) was modelled onto the Spike monomer 3 glycan site N234 (Oligomannose structure Man9 in red). Structural superposition of CLEC4G and ACE2 highlights sterical incompatibility.

Human CLEC4G and mouse Clec4g, on the other hand, were shown to have a high affinity for complex *N*-glycans terminating with GlcNAc (Pipirou et al., 2011; Powlesland et al., 2008). To further assess the detailed ligand specificity of these two lectins, we performed glycan microarray analyses comprising of 144 different glycan structures. Our analysis revealed a remarkable specificity of both, human CLEC4G and mouse Clec4g, exclusively for N-glycans with an unsubstituted GlcNAcβ-1,2Manα-1,3Man arm (Fig. 5A, Fig. S6A, B, and Table S4). In addition, our analysis revealed that mouse CD209c exhibits overlapping ligand specificities with murine Clec4g and human CLEC4G, which is in fact distinct from the known binding profile of human CD209. However, unlike Clec4g, CD209c recognized all *N*-glycan structures that displayed terminal unsubstituted GlcNAc residues, independently of the position in the glycan antennae (Fig. 5A, Fig. S6C and Table S4). The *N*-glycan at N343, located within the RBD, is the glycosylation site most abundantly decorated with terminal GlcNAc in Spike (Fig. 1C, table S2), constituting the candidate binding site for murine CD209c and murine and human CLEC4G.

The terminal GlcNAc glycans on position N343 are accessible for hCLEC4G binding on all 3 Spike monomers, but in contrast to hCD209, hCLEC4G binding interferes with the ACE2/RBD interaction (Fig. 5B, Fig. S5C, D). Since murine Clec4g and murine CD209c show strongly overlapping ligand specificities to hCLEC4G, we next modelled binding of these two lectins to the N343 glycan site. As predicted from our data, both murine Clec4g and murine CD209c indeed interfere with the ACE2/RBD interaction (Fig. S5E). Thus, whereas hCD209 is not predicted to directly affect ACE2/RBD binding, murine Clec4g, murine CD209c and human CLEC4G binding to the N343 glycan impedes Spike binding to ACE2.

### CD209c and CLEC4G block SARS-CoV-2 infection

To test our structural models experimentally, we assessed whether these lectins could interfere with Spike binding to the surface of Vero E6 cells, a frequently used SARS-CoV-2 infection model (Monteil et al., 2020). To determine this, we set-up an AFM based method, measuring spike binding activity on Vero E6 cells. Strikingly, as predicted by the structural modelling, we found that hCLEC4G, but not hCD209, significantly interfered with the binding of trimeric Spike to the Vero E6 cell surface (Fig. 6A). Similarly, mouse Clec4g, and importantly also murine CD209c, albeit to a lesser extent, interfered with Spike binding to Vero E6 cells (Fig. 6B). Finally, we tested the ability of these lectins to reduce the infectivity of SARS-CoV-2. In accordance with our model, murine Clec4g and murine CD209c significantly reduced SARS-CoV-2 infection of Vero E6 cells (Fig. 6C). Finally, hCLEC4G also significantly reduced SARS-CoV-2 infection of Vero E6 cells (Fig.6D). These data uncover that the lectins CLEC4G and CD209c can interfere with SARS-CoV-2 infections.

**Figure 6.**
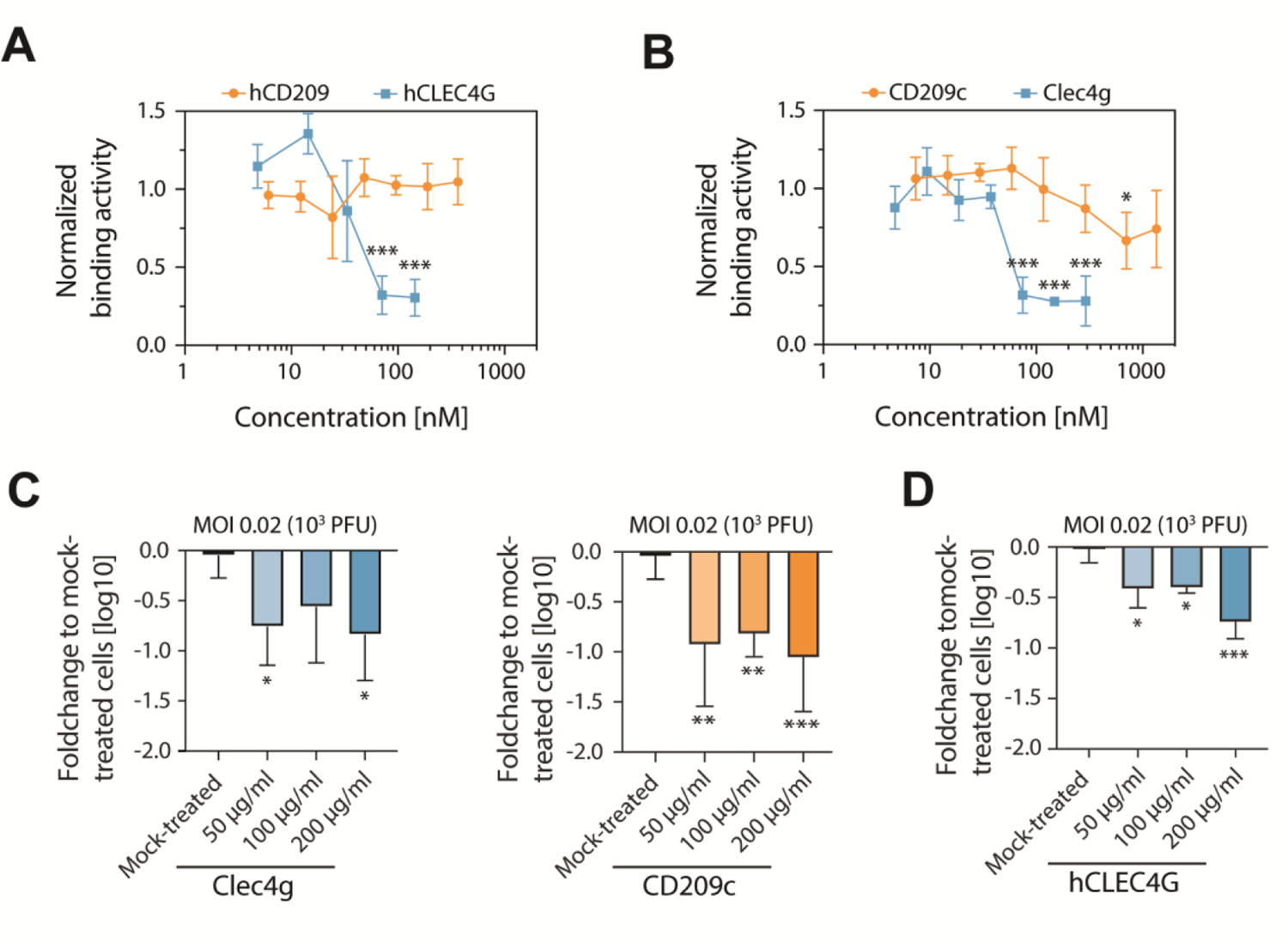
Functional determination of lectin-Spike binding in SARS-CoV-2 infections. (**A**) and (**B**) Binding activity of the full-length trimeric Spike coupled to an AFM cantilever tip to the surface of Vero E6 cells in the presence of (A) hCLEC4G and hCD209 or (B) mouse Clec4g or murine CD209c, probed at the indicated concentrations in SMFS experiments. A definite decrease in binding was observed at concentrations of 37 – 75 nM for Clec4g and 33 – 71 nM for hCLEC4G. Data are normalized to the untreated control and shown as mean ± SD (N=4). (**C**) Infectivity of mouse Clec4g or CD209c and (**D**) hCLEC4G pre-treated SARS-CoV-2 virus in Vero E6 cells. Viral RNA was measured with qRT-PCR 15 hours after infection with 0.02 MOI (10^3^ PFUs) of SARS-CoV-2. Data represent two pooled experiments for mouse lectins (N=5-6) and one experiment for hCLEC4g (N=3) and are presented as fold changes of viral loads over mock-treated controls (mean log10 values ± SD). (A) - (D) ANOVA with Dunett’s multiple comparisons test with the mock-treated group; *P<0.05; **P<0.01; ***P<0.001.

## Discussion

Here we report the results from unbiased screening of a comprehensive mammalian lectin library for potent SARS-CoV-2 Spike binding, identifying mouse CD209c and Clec4g, as well as their human homologs hCD209 and hCLEC4G, as N-glycan dependent Spike receptors. hCD209 has been identified as candidate receptor for SARS-CoV-2 by other groups, and other lectins have been also implicated in cellular interactions with Spike (Gao et al., 2020; Thépaut et al., 2020). CLEC4G has been reported to associate with SARS-CoV (Gramberg et al., 2005), but has not been implicated in SARS-CoV-2 infections. High-speed atomic force microscopy allowed us to directly observe Spike/lectin interactions in real time. Our high-speed AFM data showed that Spike/CLEC4G form more rigid clusters with lower conformational flexibility as compared to the Spike/CD209 complexes in equilibrium conditions. This is in accordance with faster association rates and shorter dissociation paths of CLEC4G as compared to CD209. The experimentally observed association of 3-4 CLEC4G molecules with one molecule of the trimeric spike indicates the formation of high affinity bonds to 1-2 glycosylation sites per monomeric subunit of the trimeric Spike.

Since glycosylation is not a template driven process, but rather depends on the coordinated action of many glycosyltransferases and glycosidases (Stanley et al., 2015), each glycosylation site can - within some boundaries - carry a range of glycans. As a consequence, the 3 monomers of Spike can harbor different glycans on the same glycosylation site. We identified N343 as the one glycosylation site that is almost exclusively covered with GlcNAc terminated glycans, the ligands of human CLEC4G and mouse Clec4g as well as, based on our new data, mouse CD209c. Given localization of N343 within the RBD, we hypothesized that CLEC4G and mCD209c binding interferes with the RBD-ACE2 interaction. Indeed, we found that murine Clec4g and human CLEC4G, acting as a multi-valent effective inhibitor (K_I_ ∼ 35 – 70 nM), can functionally impede with Spike binding to host cell membranes, thereby providing a rationale how this lectin can affect SARS-CoV-2 infections. In support of our data, it has recently been reported that a N343 glycosylation mutant exhibits reduced infectivity using pseudo-typed viruses (Li et al., 2020).

As for murine CD209c, our atomic force microscopy data indicate that CD209c engages in a larger number of interactions per trimeric Spike (∼ 5.2), presumably because mCD209c binds to a greater variety of GlcNAc-linkages than Clec4g. This more promiscuous glycan binding of mCD209c might also explain less, albeit significant, inhibition of Spike binding to VeroE6 surface as compared to human and mouse CLEC4G. Interestingly, the efficiency of inhibiting SARS-CoV-2 infection between murine and human CLEC4G and mCD209c remained comparable. As such, the multi-valent binding of CD209c may block SARS-CoV-2 infection not only through direct interference with ACE2 binding, but possibly also via affecting conformational Spike structures or proteolytic cleavage. Of note, while human and mouse CLEC4G as well as mouse CD209c can interfere with RBD-ACE2 binding, human CD209 does apparently not associate with glycans near the RBD and hence does not block Spike binding to cells. This is in agreement to its proposed high affinity oligo-mannosidic ligands, presented at N234, which is localized at a distance to the RBD-ACE2 interface.

Lectins play critical roles in multiple aspects of biology such as immune responses, vascular functions, or as endogenous receptors for various human pathogens. Hence, our library containing 143 lectins will now allow to comprehensively probe and map glycan structures on viruses, bacteria or fungi, as well as during development or on cancer cells, providing novel insights on the role of lectin-glycosylation interactions in infections, basic biology, and disease. For instance, CD209 is expressed by antigen presenting dendritic cells, as well as inflammatory macrophages (Garcia-Vallejo and van Kooyk, 2013) and is known to bind to a variety of pathogens, like HIV and Ebola, but also Mycobacterium tuberculosis or Candida albicans (Appelmelk et al., 2003). CLEC4G is strongly expressed in liver and lymph node sinusoidal endothelial cells and can also be found on stimulated dendritic cells and macrophages (Dominguez-Soto et al., 2009; Dominguez-Soto et al., 2007). CD299, one of the two homologues of mouse CD209c, which we also identified to possess Spike binding ability, is co-expressed with CLEC4G on liver and lymph node sinusoidal endothelial cells (Liu et al., 2004). Sinusoidal endothelial cells are important in the innate immune response, by acting as scavengers for pathogens as well as antigen cross-presenting cells (Knolle and Wohlleber, 2016). Thus, lectin binding to Spike might couple SARS-CoV-2 infections to antiviral immunity, which needs to be further explored. Since viral protein glycosylation depends on the glycosylation machineries of the infected cells which assemble viral particles (Watanabe et al., 2019), slight changes in glycosylation might explain differences in anti-viral immunity and possibly severity of the disease, with critical implications for vaccine designs. Moreover, Spike-binding lectins could enhance viral entry in tissues with low ACE2 expression, thus extending the organ tropism of SARS-CoV-2.

Intriguingly, all 22 *N*-glycan sites of Spike remain highly conserved among all the predominant SARS-CoV-2 variants, indicating a selection pressure to preserve these sites. Thus, the capacity of CLEC4G and mCD209c lectins to block SARS-CoV-2 viral entry holds promise for pan-variant therapeutic interventions.

## Material and Methods

### Identification of proteins containing carbohydrate recognition domains (CRDs)

Mouse lectin sequences were obtained using a domain-based approach. Briefly, proteins with a C-type lectin-like/IPR001304 domain were downloaded from InterPro 66.0 and supplemented with proteins obtained in jackhmmer searches using the PF00059.20 lectin C-type domain definition versus the mouse-specific UniProt and Ensembl databases. The collected set of candidate mouse lectins was made non-redundant using nrdb 3.0. The C-type lectin-like regions were extracted from the full-length proteins using the SMART CLECT domain definition with hmmersearch v3.1b2 and extended by 5 amino acids on both sides. To reduce redundancy, principal isoforms were selected using appris 2016_10.v24. In addition, the CRD domains for Galectins and Siglecs were added. In case a gene contained more than 1 CRD, all CRDs were cloned separately and differentiated by _1, _2, etc.

### Cloning of C-type lectin expression vectors

We used the pCAGG_00_ccb plasmid and removed the toxic ccb element by cleaving the plasmid with BsaI. Thereafter, we inserted a Fc-fusion construct, consisting of the IL2 secretion signal, followed by an EcoRV restriction site, a (GGGS)_3_ linker domain and the mouse IgG2a-Fc domain. Subsequently, each identified CRD, was cloned in-frame into the EcoRV site.

### Transfection and purification of the lectin-mIgG2a fusion proteins

The CRD containing plasmids were transfected into Freestyle^TM^ 293-F cells. Briefly, the day before transfection, 293-F cells were diluted to 0.7×10^6^ cells/ml in 30 ml Freestyle^TM^ 293-F medium and grown at 120 rpm at 37°C with 8% CO2. The next morning, 2 μl polyethylenimine (PEI) 25K (1mg/ml; Polysciences, 23966-1) per μg of plasmid DNA were mixed with pre-warmed Opti-MEM media (ThermoFisher Scientific, 31985-062) to a final volume of 950 μl in tube A. In tube B, 1 μg of DNA per ml of media was mixed with pre-warmed Opti-MEM to a final volume of 950 μl. Then, the contents of tube A and B were mixed, vortexed for 1 min and incubated at room temperature for 15 min. Thereafter, the transfection mixture was added to the cell suspension. 24h after transfection, EX-CELL 293 Serum-Free Medium (Sigma Aldrich, 14571C) was added to a final concentration of 20%. The transfected cells were grown for 120h and the supernatants, containing the secreted lectin-mIgG2a fusion proteins, harvested by centrifugation at 250g for 10 min.

Purification of the mouse lectin-mIgG2a fusion proteins was performed using Protein A agarose resin (Gold Biotechnology, P-400-5). The protein A beads were pelleted at 150g for 5 min and washed once with 1x binding buffer (0.02 M Sodium Phosphate, 0.02% sodium azide, pH=7.0), before resuspending in 1x binding buffer. Immediately preceding purification, aggregates were pelleted from the cell culture supernatant by centrifugation for 10 min at 3000g. 10x binding buffer was added to the cell culture supernatant to a final concentration of 1x as well as 4 μl of protein A beads per ml of cell culture supernatant. The bead/ supernatant mixture was incubated overnight at 4 °C. The next morning, beads were collected by centrifugation for 5 min at 150g, washed twice with 20 and 10 bead volumes of 1x binding buffer, the bead pellets transferred to a 1ml spin column (G-Biosciences, 786-811) and washed once more with 1 bead volume of 1x binding buffer. Excess buffer was removed by centrifugation at 100g for 5 sec. Lectin-mIgG2a fusion proteins were eluted from the protein A beads by resuspending the beads in 1 bead volume of Elution buffer (100mM Glycine-HCl, pH=2-3, 0.02% sodium azide). After 30 sec of incubation the elution buffer was collected into a 2 ml Eppendorf tube, containing Neutralization buffer (1M Tris, pH=9.0, 0.02% sodium azide) by centrifugation at 100g for 15 sec. Elution was performed for a total of 3 times. The 3 eluted fractions were pooled and the protein concentrations measured with the Pierce™ BCA Protein Assay Kit (ThermoFisher Scientific, 23225) using the Pierce™ Bovine Gamma Globulin Standard (ThermoFisher Scientific, 23212). To confirm the purity of the eluted lectin-IgG fusion proteins, we performed an SDS-PAGE, followed by a Coomassie staining. Briefly, 1 μg of eluted lectin-IgG fusion protein was mixed with Sample Buffer, Laemmli 2x Concentrate (Sigma-Aldrich, S3401) and heated to 95°C for 5 minutes. Thereafter, the samples were loaded onto NuPage^TM^ 4-12% Bis-Tris gels (ThermoFisher Scientific, NP0321BOX) and run in 1x MOPS buffer at 140V for 45 minutes. The gel was subsequently stained with InstantBlue^TM^ Safe Coomassie Stain (Sigma-Aldrich, ISB1L) for 1h and de-stained with distilled water. The gel picture was acquired with the ChemiDoc MP Imaging System (BioRad) in the Coomassie setting.

### Recombinant expression of SARS-CoV-2 Spike protein and the receptor binding domain (RBD)

Recombinant protein expression was performed by transient transfection of HEK293-6E cells, licensed from National Research Council (NRC) of Canada, as previously described (Durocher et al., 2002; Lobner et al., 2017). Briefly, HEK293-6E cells were cultivated in FreeStyle F17 expression medium (Thermo Fisher Scientific, A1383502) supplemented with 0.1% (v/v) Pluronic F-68 (Thermo Fisher Scientific, 24040032) and 4 mM L-glutamine (Thermo Fisher Scientific, 25030081) in shaking flasks at 37°C, 8% CO2, 80% humidity and 130 rpm in a Climo-Shaker ISF1-XC (Adolf Kühner AG). The pCAGGS vector constructs, containing either the sequence of the SARS-CoV-2 RBD (residues R319-F541) or the complete luminal domain of the Spike protein (modified by removing all arginine (R) residues from the polybasic furin cleavage site RRAR and introduction of the stabilizing point mutations K986P and V987P) were kindly provided by Florian Krammer, Icahn School of Medicine at Mount Sinai (NY, United States) (Amanat et al., 2020; Stadlbauer et al., 2020). High quality plasmid preparations for transfection were kindly provided by Rainer Hahn and Gerald Striedner (University of Natural Resources and Life Sciences, Vienna, Austria). Transient transfection of the cells was performed at a cell density of approximately 1.7×10^6^ cells/mL culture volume using a total of 1 μg of plasmid DNA and 2 μg of linear 40-kDa PEI (Polysciences, 24765-1) per mL culture volume. 48 h and 96 h after transfection, cells were supplemented with 0.5% (w/v) tryptone N1 (Organotechnie 19553) and 0.25% (w/v) D(+)-glucose (Carl Roth X997.1). Soluble proteins were harvested after 120-144 h by centrifugation (10 000 g, 15 min, 4°C).

### SARS-CoV-2 Spike mutation frequency data

Annotated SARS-CoV-2 Spike mutations were extracted from the virus repository nextstrain (https://nextstrain.org/); data are from sequences deposited until March 14^th^, 2021. The sites of N and S/T of all 22 *N*-glycan sequons (N-X-S/T) were extracted, statistically compared using two-tailed Student’s t-test and plotted with GraphPad Prism.

### Purification of recombinant trimeric Spike protein and monomeric RBD of SARS-CoV-2

For purification, the supernatants were filtered through 0.45 μm membrane filters (Merck Millipore HAWP04700), concentrated and diafiltrated against 20 mM phosphate buffer containing 500 mM NaCl and 20 mM imidazole (pH 7.4) using a Labscale TFF system equipped with a 5 kDa cut-off PelliconTM XL device (Merck Millipore, PXC005C50). His-tagged trimer Spike and monomeric RBD were captured using a 5 mL HisTrap FF crude column (Cytiva, 17528601) connected to an ÄKTA pure chromatography system (Cytiva). Bound proteins were eluted by applying a linear gradient of 20 to 500 mM imidazole over 20 column volumes. Fractions containing the protein of interest were pooled, concentrated using Vivaspin 20 Ultrafiltration Units (Sartorius, VS2011) and dialyzed against PBS (pH 7.4) at 4°C overnight using a SnakeSkin Dialysis Tubing (Thermo Fisher Scientific, 68100). The RBD was further polished by size exclusion chromatography (SEC) using a HiLoad 16/600 Superdex 200 pg column (Cytiva, 28-9893-35) equilibrated with PBS (pH 7.4). Both purified proteins were stored at −80°C until further use.

### Glycoproteomic analysis of Spike and RBD

Peptide mapping and glycoproteomic analysis of all samples were performed on in-solution proteolytic digests of the respective proteins by LC-ESI-MS(/MS). In brief, the pH of the samples was adjusted to pH 7.8 by the addition of 1 M HEPES, pH 7.8 to a final concentration of 100 mM HEPES, pH 7.8. The samples were then chemically reduced and S-alkylated using a final concentration of 10 mM dithiothreitol for 30 min at 56°C and a final concentration of 20 mM iodoacetamide for 30 min at room-temperature in the dark, respectively. To maximize protein sequence-coverage of the analysis, proteins were digested with either Trypsin (Promega), a combination of Trypsin and GluC (Promega) or Chymotrypsin (Roche). Eventually, all proteolytic digests were acidified by addition of 10% formic acid to pH 2 and directly analyzed by LC-ESI-MS(/MS) using an a capillary BioBasic C18 reversed-phase column (BioBasic-18, 150 x 0.32 mm, 5 μm, Thermo Scientific), installed in an Ultimate U3000 HPLC system (Dionex), developing a linear gradient from 95% eluent A (80 mM ammonium formate, pH 3.0, in HPLC-grade water) to 65% eluent B (80% acetonitrile in 80 mM ammonium formate, pH 3.0) over 50 min, followed by a linear gradient from 65% to 99% eluent B over 15 min, at a constant flow rate of 6 μL/min, coupled to a maXis 4G Q-TOF instrument (Bruker Daltonics; equipped with the standard ESI source). For (glyco)peptide detection and identification, the mass-spectrometer was operated in positive ion DDA mode (i.e. switching to MS/MS mode for eluting peaks), recording MS-scans in the m/z range from 150 to 2200 Th, with the 6 highest signals selected for MS/MS fragmentation. Instrument calibration was performed using a commercial ESI calibration mixture (Agilent). Site-specific profiling of protein glycosylation was performed using the dedicated Q-TOF data-analysis software packages Data Analyst (Bruker Daltonics) and Protein Scape (Bruker Daltonics), in conjunction with the MS/MS search engine MASCOT (Matrix Sciences Ltd.) for automated peptide identification.

### ELISA assays to detect lectin binding to Spike and RBD

Briefly, 50 µl of full-length Spike-H6 (4 μg/ml, purified from HEK, diluted in PBS), RBD-H6 (2 μg/ml, purified from HEK, diluted in PBS) or human recombinant soluble ACE2 (hrsACE2; 4 μg/ml, purified from CHO, diluted in PBS, see (Monteil et al., 2020)) per well were used to coat a clear flat-bottom MaxiSorp 96-well plate (Thermo Fisher Scientific, 442404) for 2h at 37°C. Thereafter, the coating solution was discarded and the plate was washed 3 times with 300 µl of wash buffer (1xTBS, 1mM CaCl_2_, 2mM MgCl_2_, 0.25% Triton X-100 (Sigma-Aldrich, T8787)). Unspecific binding was blocked with 300 µl of blocking buffer (1xTBS, 1% BSA Fraction V (Applichem, A1391,0100), 1mM CaCl_2_, 2mM MgCl_2_ and 0.1% Tween-20 (Sigma-Aldrich, P1379)) for 30 min at 37°C. After removal of the blocking solution, 50 µl of either the mouse lectin-mIgG2a (10 μg/ml, diluted in blocking buffer), human CD209-hIgG1 (R&D Systems, 161-DC-050), human CD299-hIgG1 (R&D Systems, 162-D2-050), human CLEC4G-hIgG1 (Acro Biosystems, CLG-H5250-50ug) (10 μg/ml, diluted in blocking buffer), recombinant human ACE2-mIgG1 (2 μg/ml, diluted in blocking buffer, Sino Biological, 10108-H05H) or recombinant human ACE2-hIgG1 (2 μg/ml, diluted in blocking buffer, Sino Biological, 10108-H02H) were added for 1h at room temperature. After washing for 3 times, 100 μl of 0.2 μg/ml HRP-conjugated goat anti-Mouse IgG (H+L) (Thermo Fisher Scientific, 31430) or goat anti-Human IgG (H+L) (Promega, W4031) antibodies were added for 30 min at room temperature. Subsequently, plates were washed as described above. To detect binding, 1 tablet of OPD substrate (Thermo Fisher Scientific, 34006) was dissolved in 9 ml of deionized water and 1ml of 10x Pierce™ Stable Peroxide Substrate Buffer (Thermo Fisher, Scientific, 34062). 100 μl of OPD substrate solution were added per well and incubated for 15 min at room temperature. The reaction was stopped by adding 75 μl of 2.5M sulfuric acid and absorption was read at 490 nm. Absorption was measured for each lectin-Fc fusion protein tested against full-length Spike-H6, RBD-H6 or hrsACE2 and normalized against bovine serum albumin coated control wells.

### Protein denaturation and removal of N-glycans

To denature the full-length Spike-H6, 10 mM DTT was added to 40 μg/ml of protein. The samples were incubated at 85°C for 10 min. Thereafter, the denatured proteins were diluted to 4 μg/ml with PBS and a clear flat-bottom Maxisorp 96-well plate was coated with 50 μl per well for 2h at 37°C. To remove the *N*-glycans from the full-length Spike-H6, 0.2 μg protein were denatured as above and adjusted to a final concentration of 1x Glycobuffer 2 containing 125U PNGase F per μg (NEB, P0704S) in 50 μl. After incubation for 2h at 37°C, the reaction was stopped by heat-inactivation for 10 min at 75°C. Spike proteins were then diluted to 4 μg/ml with PBS and a clear flat-bottom Maxisorp 96-well plate was coated with 50 µl per well for 2h at 37°C. ELISA protocols were performed as described above. To confirm the de-glycosylation of Spike proteins, 0.5 μg were loaded on an SDS-PAGE gel followed by a Coomassie staining.

### Surface plasmon resonance (SPR) measurements

A commercial SPR (BIAcore X, GE Healthcare, USA) was used to study the kinetics of binding and dissociation of lectin-Fc dimers to the trimeric full-length Spike in real time. Spike-H6 was immobilized on a Sensor Chip NTA (Cytiva, BR100034) via its His6-tag after washing the chip for at least 3 minutes with 350 mM EDTA and activation with a 1 min injection of 0.5 mM NiCl_2_. 50 nM Spike were injected multiple times to generate a stable surface. For the determination of kinetic and equilibrium constants, the lectin samples (murine Clec4g, murine CD209c, human CLEC4G, human CD209) were injected at different concentrations (10 to 500 nM). As the binding of the lectins is Ca^2+^ dependent, lectins were removed from the surface by washing with degassed calcium free buffer (TBS, 0.1% Tween-20, pH = 7.4). The resonance angle was recorded at a 1 Hz sampling rate in both flow cells and expressed in resonance units (1 RU = 0.0001°). The resulting experimental binding curves were fitted to the “bivalent analyte model”, assuming a two-step binding of the lectins to immobilized Spike. All evaluations were done using the BIAevaluation 3.2 software (BIAcore, GE Healthcare, USA).

### Single molecule force spectroscopy (SMFS) measurements

For single molecule force spectroscopy a maleimide-Poly(ethylene glycol) (PEG) linker was attached to 3-aminopropyltriethoxysilane (APTES)-coated atomic force microscopy (AFM) cantilevers by incubating the cantilevers for 2h in 500 µL of chloroform containing 1 mg of maleimide-PEG-N-hydroxysuccinimide (NHS) (Polypure, 21138-2790) and 30 µl of triethylamine. After 3 times washing with chloroform and drying with nitrogen gas, the cantilevers were immersed for 2h in a mixture of 100 µL of 2 mM thiol-trisNTA, 2 µL of 100 mM EDTA (pH 7.5), 5 µL of 1 M HEPES (pH 7.5), 2 µl of 100 mM tris(carboxyethyl)phosphine (TCEP) hydrochloride, and 2.5 µL of 1 M HEPES (pH 9.6) buffer, and subsequently washed with HEPES-buffered saline (HBS). Thereafter, the cantilevers were incubated for 4h in a mixture of 4 µL of 5 mM NiCl_2_ and 100 µL of 0.2 µM His-tagged Spike trimers. After washing with HBS, the cantilevers were stored in HBS at 4°C (Oh et al., 2016). For the coupling of lectins to surfaces, a maleimide-PEG linker was attached to an APTES-coated silicon nitride surface. First, 2 μl of 100 mM TCEP, 2 μl of 1M HEPES (pH 9.6), 5 μl of 1M HEPES (pH 7.5), and 2 μl of 100 mM EDTA were added to 100 μl of 200 μg/ml Protein A-Cys (pro-1992-b, Prospec, NJ, USA) in PBS. The surfaces were incubated in this solution for 2h and subsequently washed with PBS and 0.02 M sodium phosphate containing 0.02% sodium azide, pH=7.0. Finally, 100 μl of 200 μg/ml lectin-Fc fusion proteins were added to the surfaces overnight.

Force distance measurements were performed at room temperature (∼25 °C) with 0.01 N/m nominal spring constants (MSCT, Bruker) in TBS buffer containing 1 mM CaCl_2_ and 0.1 % TWEEN-20. Spring constants of AFM cantilevers were determined by measuring the thermally-driven mean-square bending of the cantilever using the equipartition theorem in an ambient environment. The deflection sensitivity was calculated from the slope of the force-distance curves recorded on a bare silicon substrate. Determined spring constants ranged from 0.008 to 0.015 N/m. Force-distance curves were acquired by recording at least 1000 curves with vertical sweep rates between 0.5 and 10 Hz at a z-range of typically 500 – 1000 nm (resulting in loading rates from 10 to 10,000 pN/s), using a commercial AFM (5500, Agilent Technologies, USA). The relationship between experimentally measured unbinding forces and parameters from the interaction potential were described by the kinetic models of Bell (Bell, 1978) and Evans and Ritchie (Evans and Ritchie, 1997). In addition, multiple parallel bond formation was calculated by the Williams model (Williams, 2003) from the parameters derived from single bond analysis. The binding probability was calculated from the number of force experiments displaying unbinding events over the total number of force experiments.

The probability density function (PDF) of unbinding force was constructed from unbinding events at the same pulling speed. For each unbinding force value, a Gaussian unitary area was computed with its center representing the unbinding force and the width (standard deviation) reflecting its measuring uncertainty (square root of the variance of the noise in the force curve). All Gaussian areas from one experimental setting were accordingly summed up and normalized with its binding activity to yield the experimental PDF of unbinding force. PDFs are equivalents of continuous histograms as shown in Fig. 3C.

### High-speed AFM (hsAFM) and data analysis

Purified SARS-CoV-2 trimeric Spike glycoproteins, murine Clec4g and CD209c and hCLEC4g and hCD209 were diluted to 20 µg/ml with imaging buffer (20mM HEPES, 1mM CaCl_2_, pH 7.4) and 1.5 µl of the protein solution was applied onto freshly cleaved mica discs with diameters of 1.5 mm. After 3 minutes, the surface was rinsed with ∼15µL imaging buffer (without drying) and the sample was mounted into the imaging chamber of the hsAFM (custom-built, RIBM, Japan). Movies were captured in imaging buffer containing 3µg/ml of either Clec4g, CD209c, hCLEC4G or hCD209. An ultra-short cantilever (USC-F1.2-k0.15 nominal spring constant 0.15 N/m, Nanoworld, Switzerland) was used and areas of 100×100nm containing single molecules were selected to capture the hsAFM movies at a scan rate of ∼150-300 ms per frame. During the acquisition of the movies, the amplitude was kept constant and set to 90-85% of the free amplitude (typically ∼3 nm). Data analysis was performed using the Gwyddion 2.55 software. Images were processed to remove background and transient noise. For volume measurements, a height threshold mask was applied over the protein structures with a minimum height of 0.25 – 0.35 nm to avoid excessive background noise in the masked area. The numbers of lectin molecules bound to the Spike trimers was calculated based on the measured mean volumes of the full-length Spike, the lectins, and the Spike-lectin complexes, averaged over the recorded time-periods.

### AFM measured Spike binding to Vero E6 cells

Vero E6 cells were grown on culture dishes using DMEM containing 10% FBS, 500 units/mL penicillin and 100 µg/mL streptomycin, at 37°C with 5% CO_2_. For AFM measurements, the cell density was adjusted to about 10-30% confluency. Before the measurements, the growth medium was exchanged to a physiological HEPES buffer containing 140 mM NaCl, 5 mM KCl, 1 mM MgCl_2_, 1 mM CaCl_2_, and 10 mM HEPES (pH 7.4). Lectins were added at the indicated concentrations. Using a full-length Spike trimer anchored to an AFM cantilever (described above), force-distance curves were recorded at room temperature on living cells with the assistance of a CCD camera for localization of the cantilever tip on selected cells. The sweep range was fixed at 3000 nm and the sweep rate was set at 1 Hz. For each cell, at least 100 force-distance cycles with 2000 data points per cycle and a typical force limit of about 30 pN were recorded.

### Glycan array analyses

The glycan microarrays were analyzed by Asparia Glycomics (San Sebastian, Spain) and prepared as described previously (Brzezicka et al., 2015). Briefly, 50 μM ligand solutions (1.25 nL, 5 drops, 250 pL drop volume) in sodium phosphate buffer (300 mM, 0.005% Tween-20, pH=8.4) were spatially arrayed employing a robotic non-contact piezoelectric spotter (SciFLEXARRAYER S11, Scienion) onto N-hydroxysuccinimide (NHS) activated glass slides (Nexterion H, Schott AG). After printing, the slides were placed in a 75 % humidity chamber for 18 hours at 25°C. The remaining NHS groups were quenched with 50 mM solution of ethanolamine in sodium borate buffer (50 mM, pH=9.0) for 1h. The slides were washed with PBST (PBS/0.05% Tween-20), PBS and water, then dried in a slide spinner and stored at −20°C until use.

Glycan microarrays were compartmentalized using Proplate® 8 wells microarray gaskets, generating 7 independent subarrays per slide. Fusion proteins were diluted at a final concentration (10 μg/mL) in binding buffer (25mM Tris, 150 mM NaCl, 4 mM CaCl2 and 0.005% Tween 20 containing 0.5 % bovine serum albumin). Lectins were applied to the microarrays and incubated at 4°C overnight with gentle shaking. The solutions were removed and arrays washed with binding buffer without BSA at room temperature. Interactions were visualized by the incubation of tetramethylrhodamine (TRITC) labelled secondary goat anti-mouse IgG antibodies (Fc specific; 1:1000 dilution in binding buffer; Life Technologies) and goat anti-Human IgG (Fc specific)-Cy3 (1:1000 dilution in binding buffer; Merck). Finally, slides were washed with binding buffer without BSA, dried in a slide spinner and scanned. Fluorescence was analyzed using an Agilent G265BA microarray scanner (Agilent Technologies). The quantification of fluorescence was done using ProScanArray Express software (Perkin Elmer) employing an adaptive circle quantification method from 50 μm (minimum spot diameter) to 300 μm (maximum spot diameter). Average RFU (relative fluorescence unit) values with local background subtraction of four spots and standard deviation of the mean were recorded using Microsoft Excel and GraphPad Prism.

### Structural modelling

Structural models of the SARS-CoV-2 Spike protein were based on the model of the fully glycosylated Spike-hACE2 complex. Experimental structures deposited in the protein databank (PDB) were used to model the complex that is formed by the binding of SARS-CoV-2 Spike and ACE2 (Walls et al., 2020; Yan et al., 2020). RBD domain in complex with ACE2 was superimposed with Spike with one open RBD domain (PDB: 6VYB) and SWISS-MODEL was used to model missing residues in Spike (Waterhouse et al., 2018) (GenBank QHD43416.1). Glycan structures in agreement with the assignments of the current work were added using the methodology outlined by Turupcu et al (Turupcu and Oostenbrink, 2017). The full model is available at the MolSSI / BioExcel COVID-19 Molecular Structure and Therapeutics Hub (https://covid.molssi.org//models/#spike-protein-in-complex-with-human-ace2-spike-spike-binding). For hCLEC4G a homology model of residues 118 – 293 was constructed using Swiss-Model (Waterhouse et al., 2018) using residues 4 – 180 of chain A of the crystal structure of the carbohydrate recognition domain of DC-SIGNR (CD299) (PDB-code 1sl6, (Guo et al., 2004)). This fragment shows a sequence identity of 36% with hCLEC4G, and the resulting model showed an overall QMEAN value of − 2.68. For mClec4g, the model consisted of residues 118 – 294, with a sequence identity of 38 % to the same template model. The resulting QMEAN value was −2.88. A calcium ion and the bound Lewis x oligosaccharide of the template were taken over into the model, indicating the location of the carbohydrate binding site. For hCD209, the crystal structure of the carbohydrate recognition domain of CD209 (DC-SIGN) complexed with Man4 (PDB code 1sl4, (Guo et al., 2004)) was used. To identify binding sites of hCLEC4G and hCD209 to the Spike-hACE2 complex, a superposition of the bound carbohydrates with the glycans on Spike was performed. For hCLEC4G, we used the complex glycans at N343 of the third monomer of Spike, with the receptor binding domain in an ‘up’ position, while N343 glycans on monomer 1 and 2 were modelled with the receptor binding domain in a ‘down’ position. For hCD209 we used the high-mannose glycan at position N234 in monomer 1-3 of Spike, respectively. These glycan structures were chosen in accordance with the full-length Spike glycoproteome.

### SARS-CoV-2 infections

Vero E6 cells were seeded in 48-well plates (5×10^4^ cells per well) (Sarstedt, 83.3923) in DMEM containing 10% FBS. 24 hours post-seeding, different concentrations of lectins were mixed with 10^3^ PFU of virus (1:1) to a final volume of 100μl per well in DMEM (resulting in a final concentration of 5% FBS). After incubation for 30 min at 37°C, Vero E6 were infected either with mixes containing lectins/SARS-CoV-2, SARS-CoV-2 alone, or mock infected. 15 hours post-infection, supernatants were removed, cells were washed 3 times with PBS and then lysed using Trizol Reagent (Thermo Fisher Scientific, 15596026). The qRT-PCR for the detection of viral RNA was performed as previously described (Monteil et al., 2020). Briefly, RNA was extracted using the Direct-zol RNA MiniPrep kit (Zymo Research, R2051). The qRT-PCR was performed for the SARS-CoV-2 E gene and RNase P was used as an endogenous gene control to normalize viral RNA levels to the cell number. Lectins were independently tested for cellular toxicity in an ATP-dependent assay (Cell-Titer Glo, Promega) and cells found to exhibit >80% viability up to a concentration of 200 μg/ml (data not shown).

The following PCR Primers were used: SARS-CoV2 E-gene:

Forward primer: 5’-ACAGGTACGTTAATAGTTAATAGCGT-3’

Reverse primer: 5’-ATATTGCAGCAGTACGCACACA-3’

Probe: FAM-ACACTAGCCATCCTTACTGCGCTTCG-QSY

RNase P:

Forward primer: 5’-AGATTTGGACCTGCGAGCG-3’

Reverse primer 5’-GAGCGGCTGTCTCCACAAGT-3’

Probe: FAM-TTCTGACCTGAAGGCTCTGCGCG-MGB

## Acknowledgements

We thank all members of the Penninger laboratory for helpful discussions and technical support. Moreover, we thank all members of the Molecular Biology Service and the VBCF Protein Technologies Facility. We thank Florian Krammer (Icahn School of Medicine at Mount Sinai, NY, United States) for providing the constructs used for production of recombinant Spike and RBD. Purified Spike and RBD as well as transfection-grade pCAGGS plasmids were obtained from the reagent repository of the BOKU COVID-19 Initiative. The authors thank Daniel Maresch (BOKU Core Facility Mass Spectrometry) for assistance with glycan analysis. Fig. 1A, 3A and S1A created with BioRender.com. J.M.P. and the research leading to these results has received funding from the T. von Zastrow foundation, the FWF Wittgenstein award (Z 271-B19), the Austrian Academy of Sciences, the Innovative Medicines Initiative 2 Joint Undertaking (JU) under grant agreement No 101005026, and the Canada 150 Research Chairs Program F18-01336 as well as the Canadian Institutes of Health Research COVID-19 grants F20-02343 and F20-02015. D.H. is supported by the T. von Zastrow foundation, S.M. is funded by the European Union’s Horizon 2020 research and innovation programme under the Marie Sklodowska-Curie grant agreement No 841319. We further acknowledge financial support from the Austrian National Foundation for Research, Technology, and Development and Research Department of the State of Upper Austria (Y.J.O), from the FWF projects V584 (Y.J.O), P31599 (R.Z.), I3173 (P.H., L.H.), the WWTF grant LS19-029 (P.H, D.C.) and grant COV20-015 (C.O.), the ÖAW fellowship STIP13202002 (L.H.), and the European Union’s Horizon research and innovation programme (H2020-MSCA-ITN-2016) under the Marie Sklodowska-Curie grant agreement No. 721874 (P.H., D.C.). A.M. and V.M. have received funding from the Innovative Medicines Initiative 2 JU under grant agreement no. 101005026. JU receives support from the European Union’s Horizon 2020 research and innovation programme and EFPIA.

## Author contributions

D.H., S.M. and J.M.P. conceived and designed the study; Y.J.O. and P.H. conceived and coordinated the SPR and AFM studies; Y.J.O. performed the lectin force spectroscopy measurements and data analysis; R.Z. performed the binding activity measurements and data analysis; D.C. performed the high-speed AFM measurements and data analysis; L.H. performed the SPR measurements and data analysis; V.M. and A.M. designed, performed and analyzed the SARS-CoV-2 infection experiments; E.L. and L.M. set-up and performed the purification of the Spike and RBD; C.G.G., F.A. and J.S. designed, performed and analyzed the glycosylation of the Spike and RBD; D.H., G.W., M.N., A.C. and M.T. designed, set-up and purified the lectin-Fc proteins; D.H., A.H. and S.M. performed all ELISA experiments; C.O. performed the modelling; Y.J.O., R.Z., D.C. and L.H. wrote the original SPR and AFM part of this manuscript with guidance and edits from P.H.; D.H., S.M. and J.M.P. wrote the manuscript. All authors read and reviewed of the manuscript.

## Conflict of interest

A patent is being prepared to use CLEC4G as potential therapy for COVID-19. J.M.P. is shareholder and board member of Apeiron Biologics that is developing soluble ACE2 for COVID-19 therapy.

## Supplementary Materials

**Figure S1.**
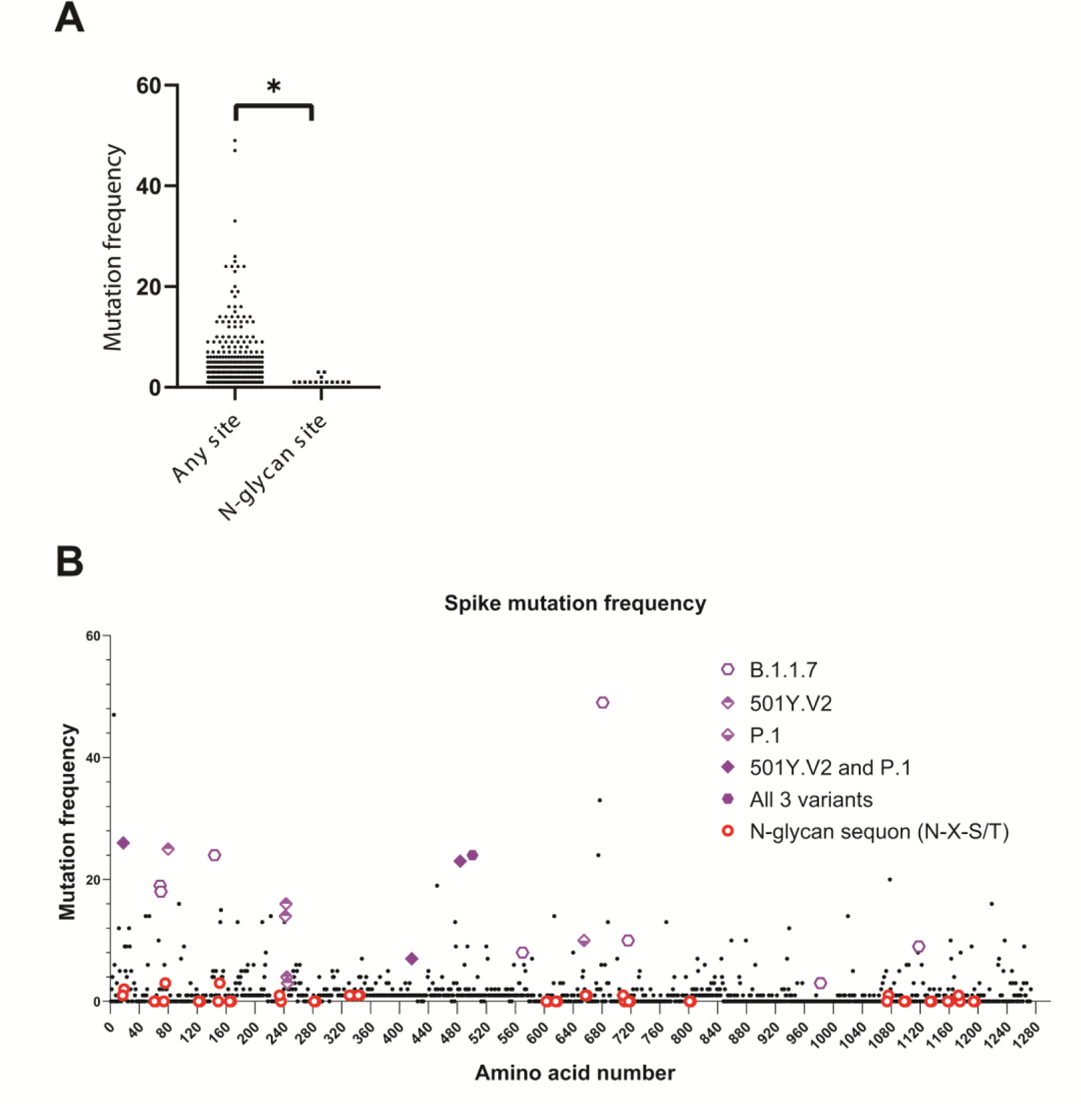
Mutation frequency of *N*-glycan sites on SARS-CoV-2 Spike. (A) Among the 1273 amino acids of Spike the frequency of mutational amino acid conversion within N-glycan sequons is plotted again all other sites. **(B)** The mutation frequency of all 1273 amino acids of Spike is shown. *N*-glycan sequons as well as mutations harbored by the new variants B.1.1.7, 501Y.V2 and P.1 are highlighted. (A) Two-tailed Student’s T-test, *p < 0.05.

**Figure S2.**
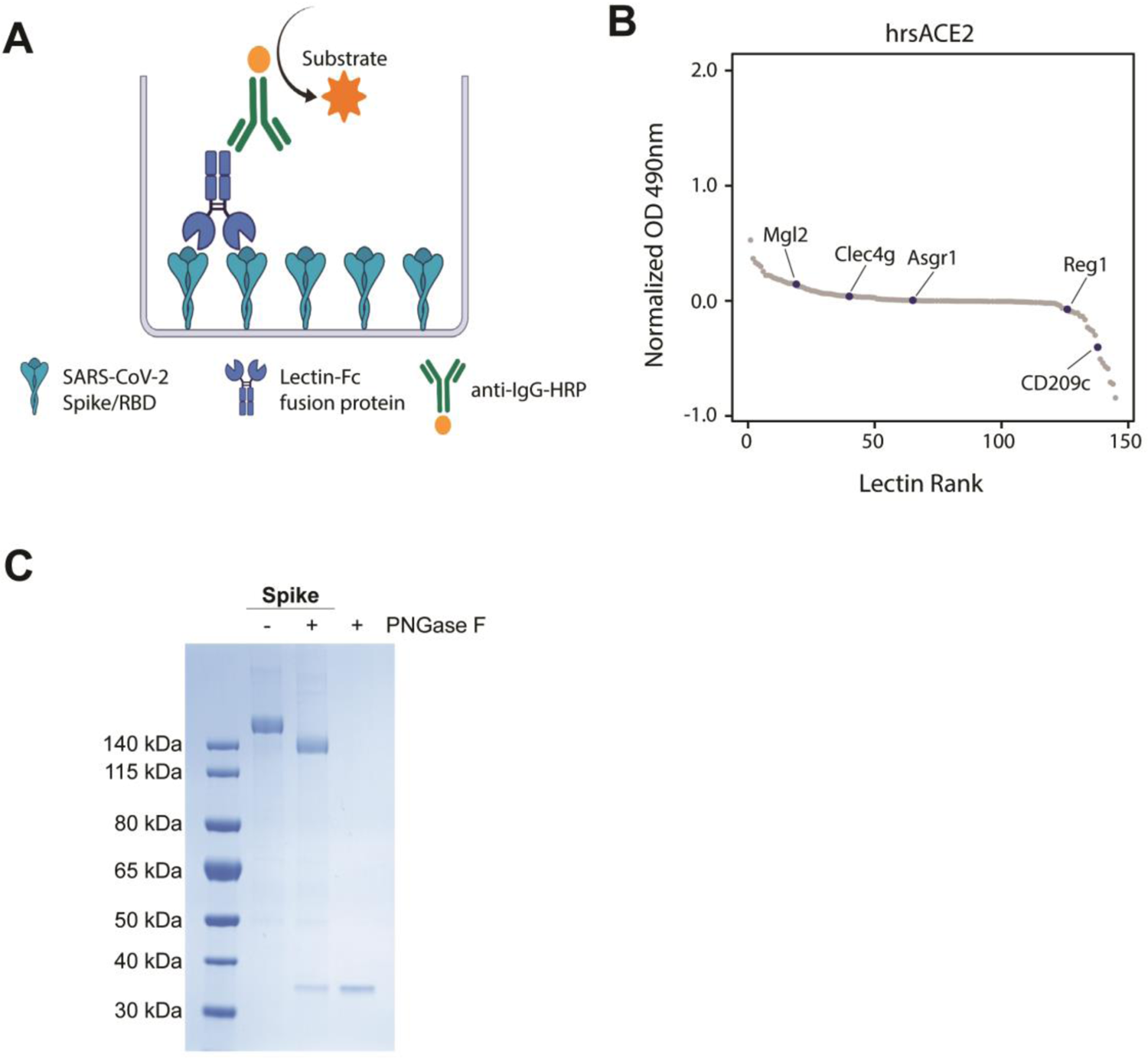
ELISA assays to detect lectin binding. (**A**) Schematic representation of the ELISA protocol, consisting of coating with trimeric full-length Spike or the monomeric receptor binding domain (RBD) followed by sequential incubation with lectin-Fc fusion proteins and secondary anti-IgG-HRP antibodies. The binding of lectin-Fc fusion proteins was quantified by peroxidase-dependent substrate conversion, measured by optical density (OD) at 490nm and normalized against a BSA control. (**B**) ELISA screen of the lectin-Fc library against human recombinant soluble ACE2 (hrsACE2). Results are shown as mean OD values of 2 replicates normalized against a BSA control and ranked by value. (**C**) SDS-Page of full-length Spike de-*N*-glycosylated with PNGase F and stained with Coomassie blue. A PNGase F control was added to display the size of the PNGase F protein.

**Figure S3.**
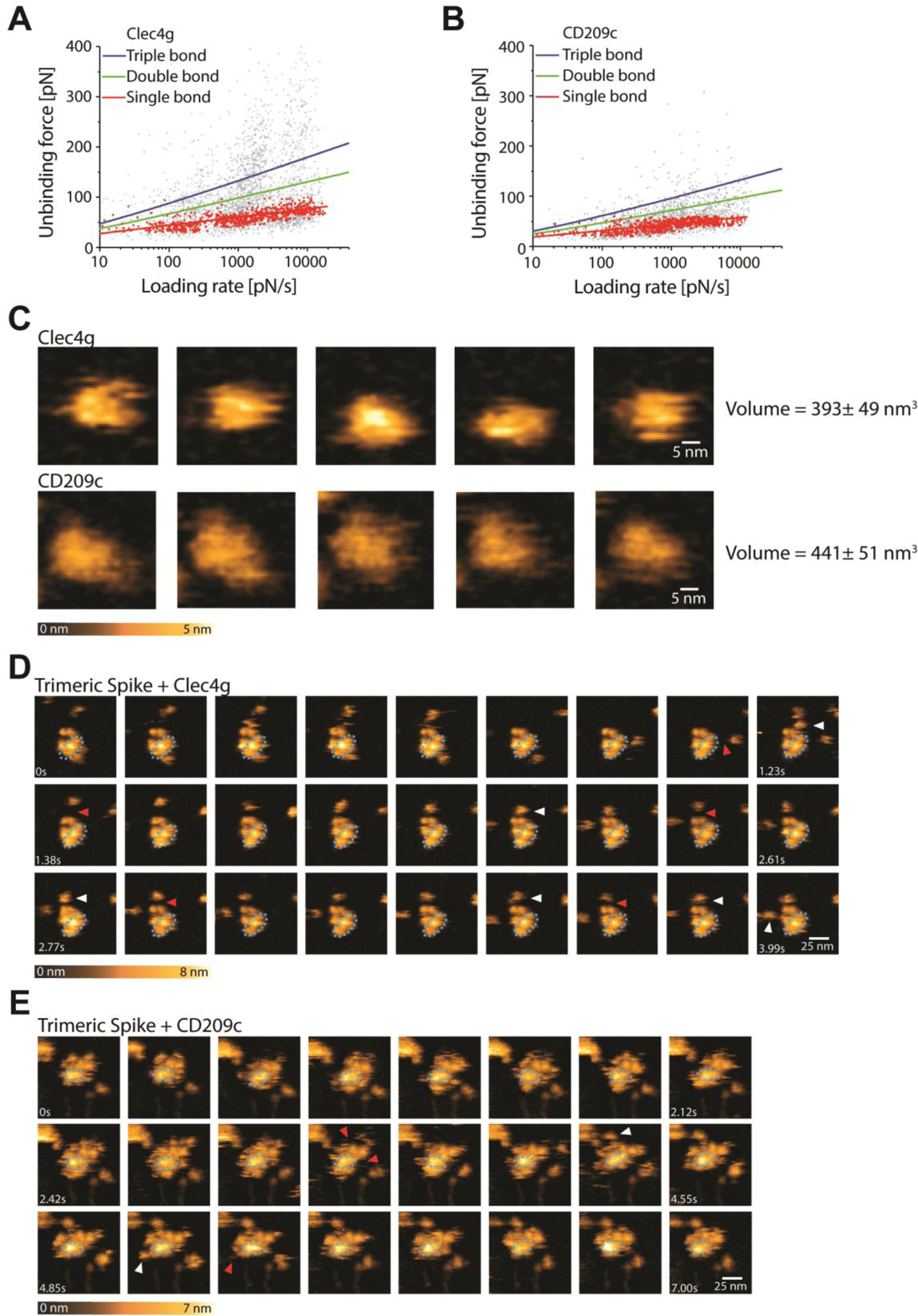
Single molecule atomic force microscopy of a single trimeric Spike binding to murine Clec4g or CD209c. (**A**) and (**B**) Unbinding forces versus loading rates for trimeric Spike dissociating from (A) Clec4g-Fc or (B) CD209c-Fc. Unbinding forces were determined from the magnitude of the vertical jumps measured during pulling of the cantilever (Fig. 3B) and individually plotted versus the respective force loading rates (equal to the pulling speed times effective spring constant) to decipher the dissociation dynamics (Table 1). A well-defined single-bond behavior of a unique monovalent bond was found (red dots) that, in line with Evans’s single energy barrier model, yielded a linear rise of the unbinding force with respect to a logarithmically increasing loading rates for both (A) Clec4g and (B) CD209c. Double (green) and triple (blue) bond behaviors were calculated according to the Markov binding model using parameters derived from the single barrier model. Unbinding force values scattered between single and triple bond strengths, indicating that interactions with various glycosylation sites with different binding strengths. pN=picoNewton, pN/s = picoNewton per second. (**C**)-(**E**) High speed AFM of a single trimeric Spike visualizing the real-time interaction dynamics with lectins. (C) 5 frames of Clec4g or CD209c alone imaged on mica. (D) Sequential movie frames of trimeric Spike/Clec4g complexes, acquired at a rate of 153.6 ms/frame, corresponding to Fig. 3C. (E) Sequential movie frames of trimeric Spike/CD209c complexes, acquired at 303 ms/frame, corresponding to Fig. 3C. White arrows point to lectins associating with the Spike trimer body. Red arrows indicate dissociation of lectins from the Spike trimer, highlighting positions where the lectin was bound in the previous frame. Blue dotted ellipses display low mobility regions. Color schemes indicate height in nanometers (nm).

**Figure S4.**
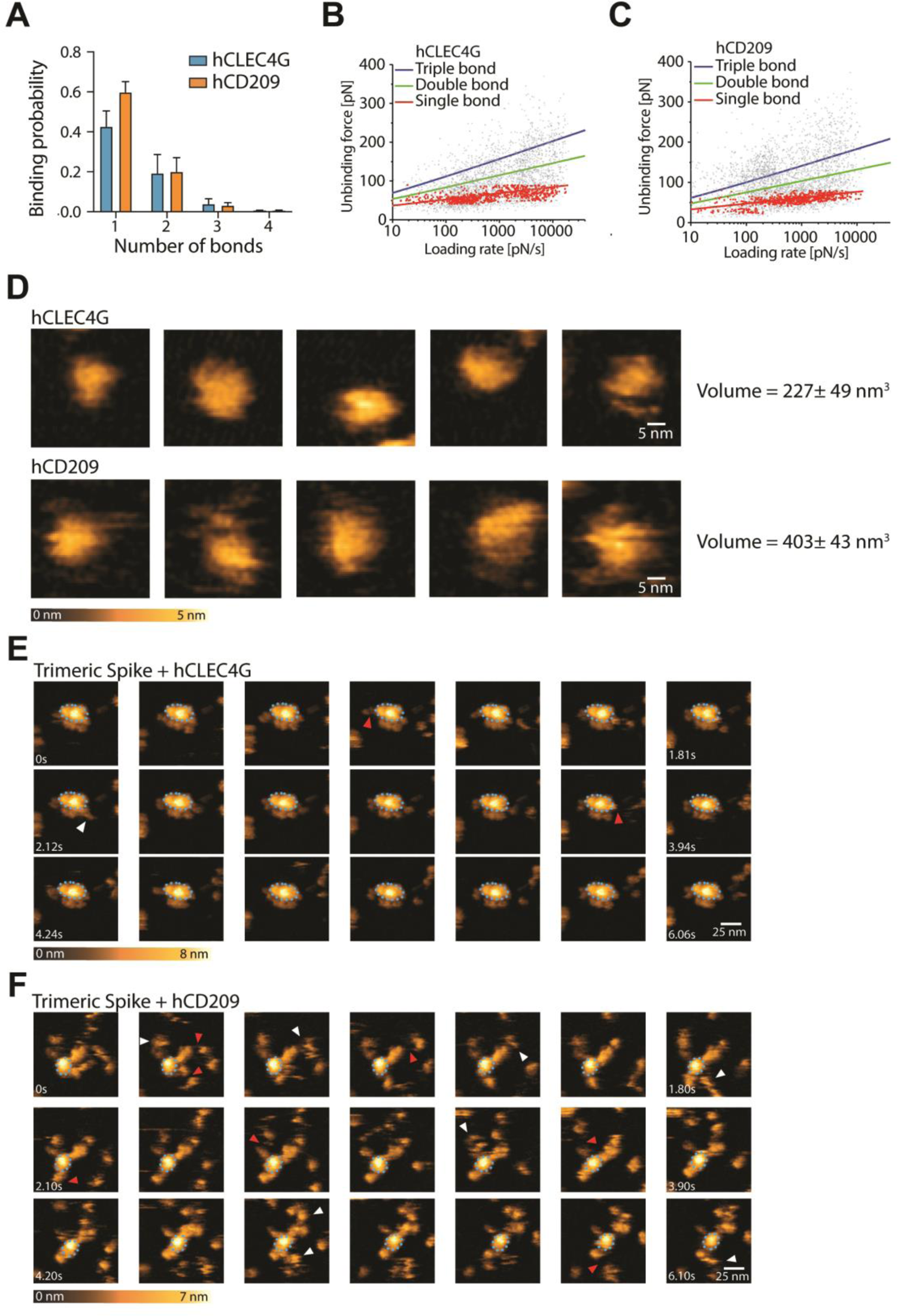
Single molecule atomic force microscopy of a single trimeric Spike binding to human CLEC4g or CD209. (**A**) Single molecule force spectroscopy (SMFS) to determine the binding probability for trimeric Spike to mica coated hCLEC4G and hCD209. Data are shown as mean binding probabilities ± SD of single, double, triple or quadruple bonds (N=2). (**B**) and (**C**) Unbinding forces versus loading rates for a single trimeric Spike dissociating from (B) hCLEC4G or (C) hCD209. Unbinding forces were determined from the magnitude of the vertical jumps measured during pulling (Fig. 3B) and individually plotted vs. their force loading rates (equal to the pulling speed times effective spring constant) to assess the dissociation dynamics (Table 1). Single bond interactions (red dots) were fitted using the Bell-Evans single barrier model (red line). A well-defined single-bond behavior of a unique monovalent bond was found (red dots) that, in line with Evans’s single energy barrier model, yielded a linear rise of the unbinding force with respect to a logarithmically increasing loading rate for both (B) hCLEC4g and (C) hCD209. Double (green) and triple (blue) bond behaviors were calculated according to the Markov binding model using parameters derived from the single barrier model. Unbinding force values scattered between single and triple bond strengths, indicating that they arise from multiple interactions with various glycosylation sites. pN=picoNewton, pN/s = picoNewton per second. (**D**)-(**F**) High speed AFM of a single trimeric Spike visualizing the real-time interaction dynamics with lectins. (D) 5 frames of hCLEC4g or hCD209 alone imaged on mica. (E) Sequential movie frames of trimeric Spike/hCLEC4g complexes, acquired at a rate of 303 ms/frame, corresponding to Fig. 4E. (F) Sequential movie frames of trimeric Spike/hCD209 complexes, acquired at 153.6 ms/frame, corresponding to Fig. 4E. White arrows point to lectins associating with the Spike trimer body. Red arrows indicate dissociation of lectins from the Spike trimer, highlighting positions where the lectin was bound in the previous frame. The blue dotted ellipses display low mobility regions. Color schemes indicate height in nanometers (nm).

**Figure S5.**
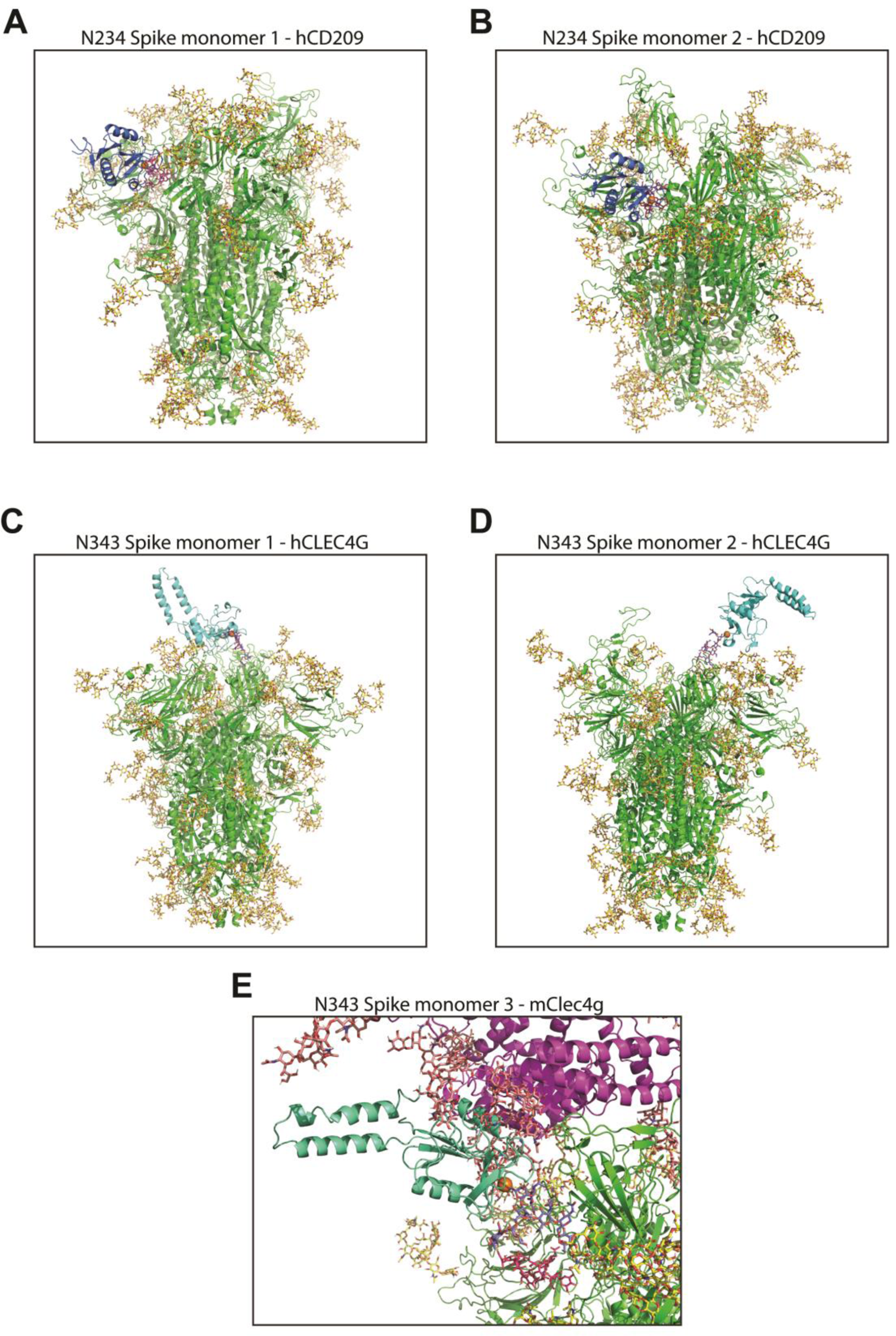
Structural modelling of lectin-Spike interactions. (**A**) and (**B**) 3D structural modelling of glycosylated trimeric Spike (green with glycans in yellow) interacting with the CRD of hCD209 (dark blue with Ca^2+^ in orange). The model shows the (A) Spike monomer 1 and (B) Spike monomer 2 glycan site N234 (Oligomannose structure Man9 in red) bound to hCD209. (**C**) and (**D**) 3D structural modelling of glycosylated trimeric Spike (green with glycans in yellow) interacting with the CRD of hCLEC4g (cyan with Ca^2+^ in orange). The model shows Spike (A) monomer 1 and (B) monomer 2 glycan site N343 (complex type glycan with terminal GlcNAc in purple-blue) bound to hCLEC4g. (**E**) 3D structural modelling of glycosylated trimeric Spike (green with glycans in yellow) interacting with glycosylated human ACE2 (purple with glycans in salmon). The CRD of mClec4g (cyan with Ca^2+^ in orange) was modelled onto Spike monomer 3 glycan site N343 (complex type glycan with terminal GlcNAc in purple-blue). Structural superposition of mClec4g and ACE2 highlights steric incompatibility.

**Figure S6.**
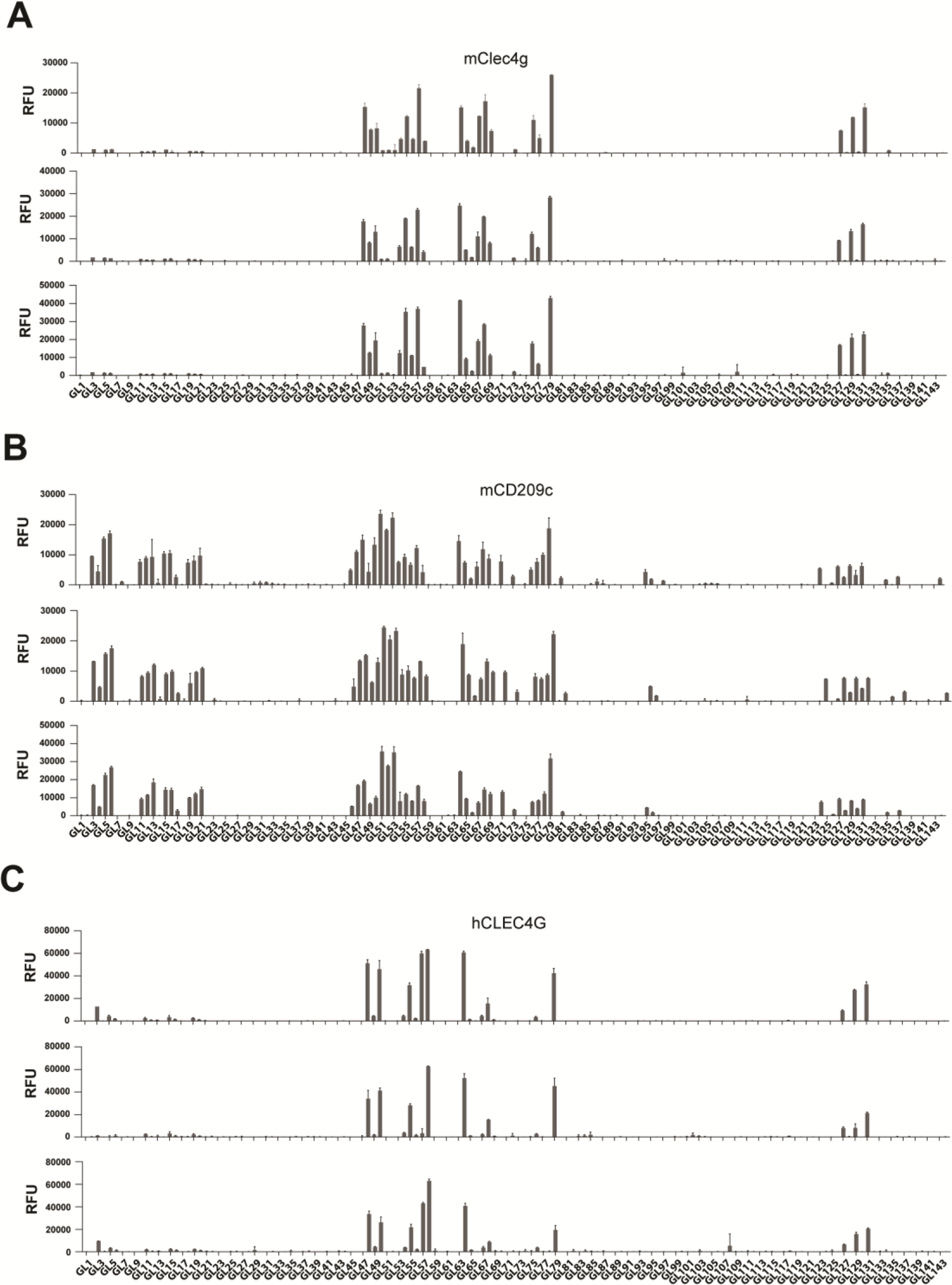
Glycan microarray results. Binding of mClec4g, mCD209c and hCLEC4G to glycans spotted on the microarray. Primary data from three independent microarray measurements are shown. Each histogram shows the average values of relative fluorescence units (RFU) +/- standard deviation. Glycan structures of GL1 to GL144 is represented in Fig S7.

**Figure S7.**
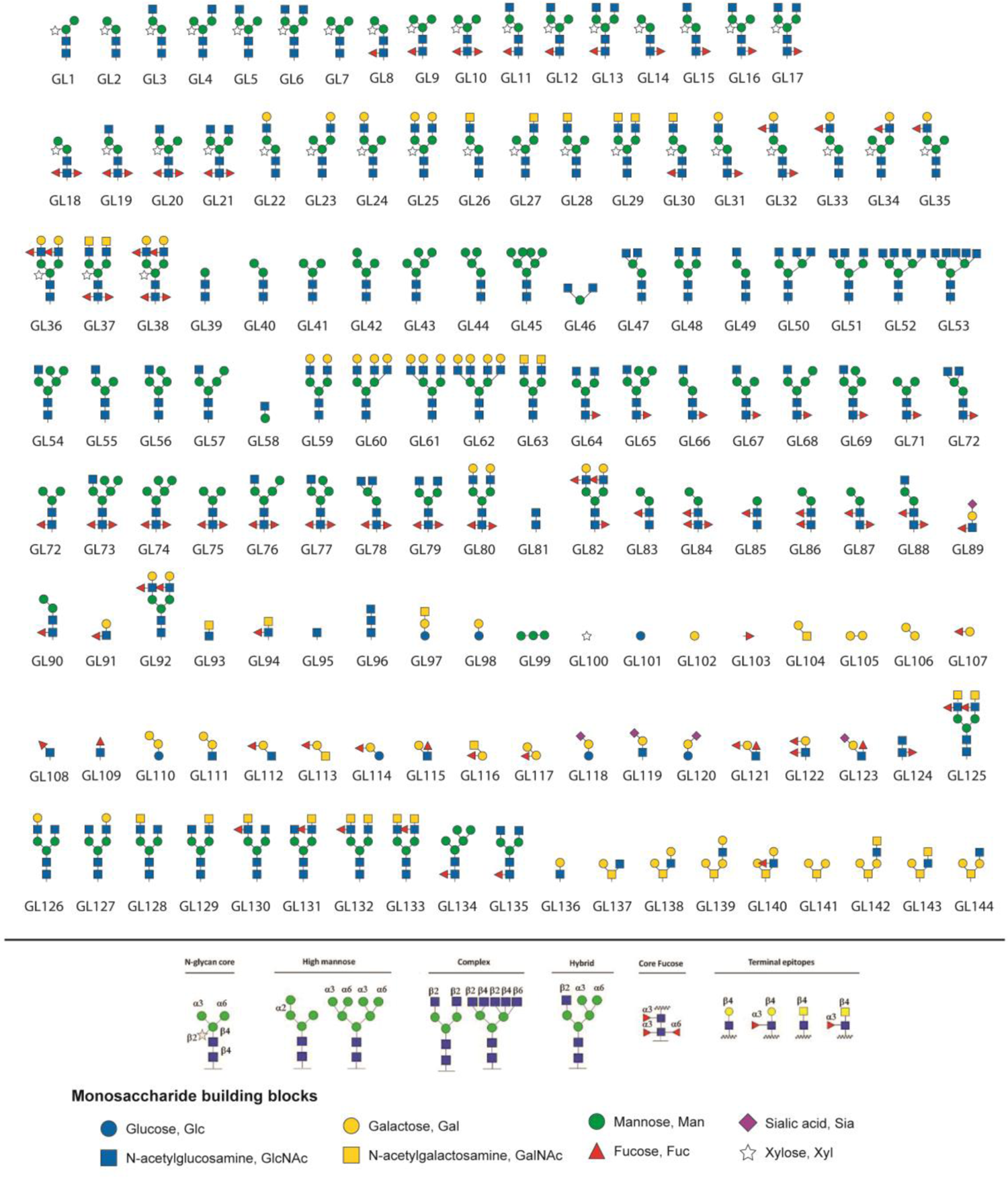
Representation of the Glycan microarray.

**Table S1. Overview of carbohydrate recognition domains (CRDs) used for the lectin library** This table presents a list of CRDs expressed and purified as Fc-fusion proteins for the lectin library. Information displayed are the lectin name, the family and in the case of C-type lectins, the group and group name the CRD belongs to. CRDs from lectin that contain several CRDs are distinguished by suffix numbers. CTL = C-type lectin

**Table S2. Glycosylation of SARS-CoV-2 Spike and RBD** Relative abundance of all measured glycans in % of all glycans present at each position. Glycans are grouped in families consisting of designated glycan features.

**Table S3. ELISA screen of the lectin-Fc library against SARS-CoV-2 Spike, RBD and hrsACE2** Results from the ELISA screens of the lectin-Fc library against indicated targets. Data displayed per lectin is mean and standard deviation (SD) (n=2). SD=NA indicates that mean was calculated from a single replicate only.

**Table S4. Glycan microarray replicates of mClec4g, mCD209c, and hCLEC4G.** Represented is the average (Avg) and standard deviation (SD) of the relative fluorescence units (RFU) measured among 4 replicated spots (n = 4). The glycan structures of GL1 to GL144 are illustrated in Fig. S7.

---

**Movie S1.**

High speed AFM of single trimeric Spike visualizing the real-time interaction dynamics with mClec4g acquired at a rate of 153.6 ms/frame.

**Movie S2.**

High speed AFM of single trimeric Spike visualizing the real-time interaction dynamics with mCD209c acquired at a rate of 303 ms/frame.

**Movie S3.**

High speed AFM of single trimeric Spike visualizing the real-time interaction dynamics with hCLEC4G acquired at a rate of 303 ms/frame.

**Movie S4.**

High speed AFM of single trimeric Spike visualizing the real-time interaction dynamics with hCD209 acquired at a rate of 303 ms/frame.

